# Retrotransposon Activation in the Aged and Alzheimer’s Disease Brain Examined by Nanopore Long-read DNA Sequencing

**DOI:** 10.64898/2025.12.12.693943

**Authors:** Maxfield M.G. Kelsey, Anjalika Chongtham, John LaCava, Martin S. Taylor, Jef D. Boeke, Fred H. Gage, Andrei Seluanov, Vera Gorbunova, Ana C. Pereira, John M. Sedivy

## Abstract

**Background:** Cellular defenses against retrotransposable elements (RTEs) weaken with age and RTEs have been reported to contribute to Alzheimer’s disease (AD) pathogenesis by promoting neuroinflammation. The mechanisms implicated include DNA damage promoted by retrotransposition and interferon system activation by RTE-derived cDNA intermediates. LINE-1 (L1) retrotransposons are of particular interest because they are the only autonomously active RTEs in the human genome.

**Results:** To investigate L1 activation and retrotransposition in AD, we performed Nanopore long-read DNA sequencing on six late-onset AD (LOAD) and six age-matched control human prefrontal cortex (PFC) samples. We developed and validated a stringent RTE insertion calling pipeline and identified two high-confidence somatic insertions, one AluY and one L1HS. We estimate that ∼1% of cells in the aged PFC have a somatic RTE insertion. AD samples were hypomethylated, and genome-wide analysis of differentially methylated regions (DMRs) supports a process of epigenetic drift in AD. DMR-associated gene sets primarily related to brain function and inflammation. To investigate L1 activation we used CpG methylation as a proxy for L1 expression. We observed decreased methylation at young L1 elements. While most reads overlapping the L1HS promoter were highly methylated (>80% methylated), 7% were <50% methylated, 1% were <25%, and the highly demethylated read fraction increased in AD. L1HS 5’ UTR methylation was strongly correlated with RNA expression.

**Conclusions:** CpG methylation-mediated repression of young RTEs is compromised in old age - our findings indicate that this is further exacerbated in AD. Amid these failing defenses, we report somatic retrotransposition events in the aging and demented brain.

## Background

Alzheimer’s disease (AD) is a form of dementia characterized by cognitive decline, neuron loss, and the deposit of amyloid beta plaques and neurofibrillary-Tau tangles[1]. The etiology of AD pathogenesis is complex, and neuroinflammation is increasingly appreciated as playing a significant role in disease development[2]. The reasons for, and processes underpinning, chronic neuroinflammation remain incompletely understood[3].

The leading risk factor for AD is age[4]. Among the many cellular processes affected by aging, the disruption of gene regulatory networks and epigenomic integrity has attracted considerable attention[5]. The capacity to restore the epigenetic landscape following perturbations such as DNA damage declines with age[6], leading to compromise of finely tuned transcriptional networks. Heterochromatic regions can become derepressed, leading to the aberrant expression of retrotransposable elements (RTEs)[7–9]. These elements constitute a reservoir of immunostimulatory pathogen-associated molecular patterns (PAMPs): RTE transcripts can form dsRNA and/or present abundant CpG motifs; RTE-encoded reverse transcriptase can also lead to the production of cytoplasmic RNA:DNA hybrids and/or dsDNAs[10, 11]. Together, these findings have prompted the hypothesis that the derepression of RTEs in the aged brain serves as a cell-intrinsic stimulus driving neuroinflammation and promoting neurodegeneration.

The defining feature of the RTE lifecycle is a reliance on RNA intermediates that must be reverse transcribed prior to or during integration into the genome. While these elements account for roughly 40% of the human genome, the vast majority are no longer capable of retrotransposition[12]. Most have lost their promoters; others are full-length but have inactivating mutations in their protein-coding regions. Only the long-<u>in</u>terspersed <u>e</u>lement-1 (LINE-1; L1) family remains autonomously competent for retrotransposition in humans. The evolutionarily youngest L1 subfamily, L1 *Homo sapiens* (L1HS), comprises approximately 6,000 loci, ∼300 of which are full-length (>6 kb). Only ∼150 of the full-length elements retain the full complement of intact open reading frames required for retrotransposition. Given their evolutionarily recent activity in human genomes, these young elements are highly polymorphic and many L1HS insertions in an individual may be absent from the human reference genome. These non-reference elements, as evidenced by their recent activity in the human lineage, comprise some of the most active elements in our genome.

When expressed, young L1 elements can promote a range of biological events. One class of events is genotoxic stress. The L1 mechanism of retrotransposition, target-primed reverse transcription (TPRT), involves a series of nicks to genomic DNA that can lead to double-stranded (dsDNA) breaks in certain cell line models[13]. Canonically, nicking of the genomic DNA and reverse transcription of the L1 RNA is carried out by the L1-encoded ORF2 protein; additionally, ORF2 protein is necessary for the mobilization of <u>s</u>hort-<u>in</u>terspersed <u>e</u>lements (SINEs) of the Alu and SVA families. Thus, ORF2p-driven *de novo* insertions may disrupt gene function or regulation.

Another threat comes from the high levels of homology between RTE elements, which can promote non-allelic homologous recombination (NAHR)[14, 15]. It has more recently been appreciated that L1 activity engages cellular innate anti-viral defenses in somatic cells. L1 reverse transcribed cDNA registers as “non-self”, triggering cGAS-STING activation and downstream interferon signaling[16].

As a consequence of these threats, expression of retrotransposons, and in particular L1HS, is highly regulated. The first line of defense against L1 expression is DNA CpG methylation in the L1 5’ untranslated region (5’ UTR), a region that acts as an internal promoter[17–19]. Additionally, repressive histone marks such as H3K9me3 and H3K27me3 help maintain a condensed chromatin conformation at the L1 promoter[20]. Given these defenses, young L1s are not expressed in healthy cells and tissues. However, in aged and diseased cells, these defenses can weaken and enable L1 derepression[21, 22]. The extent to which this contributes to the pathogenesis of neurodegenerative conditions is an area of active investigation.

Alterations to the AD epigenome have been widely reported. Immunohistochemical studies have reported conflicting findings regarding global changes in 5-methylcytosine (5mC), with different groups finding increased, decreased, or unchanged global levels[23–25]. Sequencing based technologies including RNA-seq[26], ATAC-seq[26] and Hi-C[27] paint a more consistent picture of epigenomic erosion in AD. Wang et al. reported a loss of defined 3D genome structure in LOAD neurons[27] whereas Xiong et al. found that repressive chromatin regions become more accessible while open chromatin regions are condensed[26]. The heterochromatin loss observed in AD neurons coincides with a reactivation of normally silenced genes[28]. Further supporting a connection between AD and L1 derepression, pathogenic Tau has been shown to promote global chromatin relaxation as well as RTE expression[28–30].

To comprehensively and directly compare the DNA methylation status of L1 elements genome-wide in the aged and AD brain, we used Nanopore long-read DNA whole genome sequencing (WGS). This technology provides a number of advantages important to the study of repetitive DNA elements, enabling us to: (i) accurately map reads to repetitive genomic regions, (ii) infer, and map to, specific non-reference L1 germline/somatic insertions in the specific genome under study, and (iii) profile CpG methylation. Additionally, while this is a bulk DNA sequencing technique, long reads allow us to examine the methylation status of groups of adjacent CpGs derived from a single cell. This allows us to distinguish scenarios in which (1) a majority of cells are marginally demethylated at many L1 loci from those in which either (2a) a small percentage of cells are profoundly demethylated or (2b) many cells stochastically lose substantial amounts of methylation at unique loci. While scenarios (2a) and (2b) cannot be distinguished from one another, both are more biologically significant than is scenario (1) as far as their likely consequences on L1 expression.

## Results

### Analysis of germline and somatic RTE insertions by Nanopore DNA-seq

To characterize the RTE insertion profile and methylome of cognitively normal versus AD individuals, we performed Nanopore sequencing on DNA isolated from six AD and six control (CTRL) postmortem prefrontal cortex (PFC) samples. Samples were assigned to AD versus CTRL groups based on clinical tests of cognition and were matched for age and sex (Methods, Table S1). CTRL sample Braak stages ranged from I–IV and AD samples ranged from V–VI. Each sample was sequenced on its own PromethION flow cell. Sequencing coverage was high, surpassing 20x haploid genome coverage for all samples, averaging 34x (Table S1). Read length, as measured by N50 value, was also high for all samples (>5.5 kb).

We identified non-reference RTE insertions relative to the T2T-HS1 human reference genome using the TLDR program[31]. We first examined non-reference germline insertions, which are supported by roughly half (heterozygous insert) or all (homozygous insert) reads at a given insert site. This subset of insertions is expected to include both very recently transposed germline insertions “private” to the individual being sequenced or in that individual’s immediate ancestors), and relatively older insertions that are commonly found in the human population but are absent from the human genome reference sequence used (T2T-HS1). Among the youngest RTE subfamilies, members of which are either capable of retrotransposition or were in our recent evolutionary past, we identified comparable numbers of insertions across samples (Fig. 1A).

**Fig. 1.**
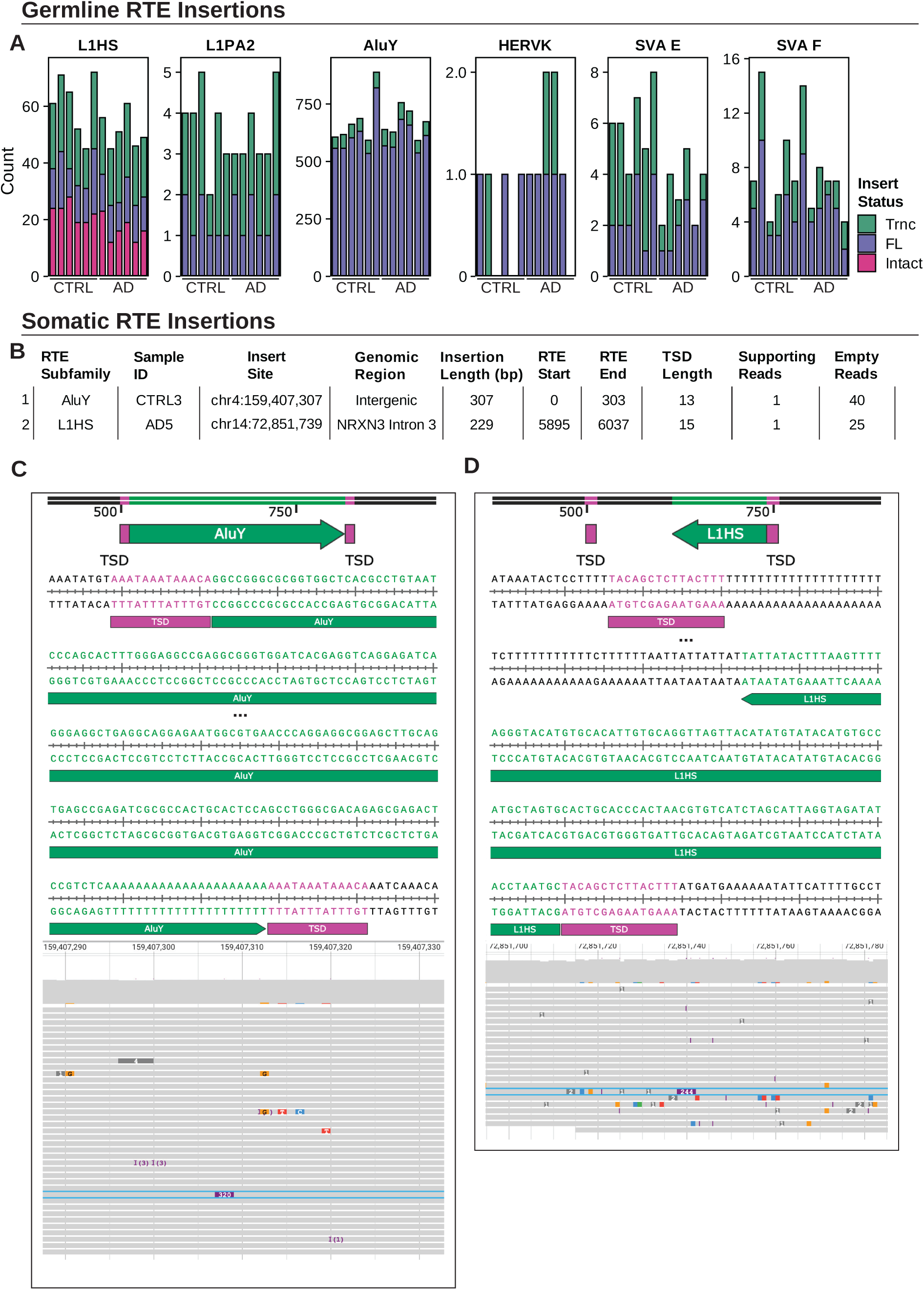
Nanopore DNA-Seq reveals non-reference germline and somatic RTE insertions. RTE insertions were called relative to the T2T-HS1 human reference genome using TLDR. Putative germline and somatic insertions were rigorously filtered (see Methods). (**A**) Polymorphic germline insertion counts for L1HS, L1PA2, AluY, HERVK, SVA E, and SVA F subfamilies. Insertions are colored by length and open reading frame (ORF) intactness. FL: full-length; Intact: intact ORFs; Trnc: truncated. (**B**) Table summarizing all observed putative somatic RTE insertion events. (**C,D**) Diagram of somatic insertion (Insert #1 (**C**) and #2 (**D**) (top). Insertion sequences were annotated by RepeatMasker for repetitive element content and by TLDR for target site duplications (TSDs). SnapGene representation of the annotated insert sequence (middle). A portion of the insert is omitted due to space constraints (denoted by ellipses). Genome browser view of all sequencing reads (unfiltered set of alignments) at the insert site (bottom). The insert spanning read is outlined in blue. Departures from the reference sequence are colored, and insertions are denoted by colored boxes indicating insert length.

AluY insertions were most common, averaging 672 insertions per sample, followed by L1HS, 56.2 insertions, SVA F, 7.8 insertions, SVA E, 4.7 insertions, L1PA2, 3.6, and HERVK, 1.2 insertions. Roughly half of the L1HS insertions were full-length, and a majority of the FL insertions had intact open reading frames for both ORF1 and ORF2 proteins and hence may be capable of retrotransposition. The small number of L1PA2 insertions were all truncated. The vast majority of RTE insertions were either intergenic or intronic (Fig. S1A). Most insertions were heterozygous, consistent with their non-reference status, though roughly a quarter of insertions were homozygous (Fig. S1B). Of the Alu insertions we detected, 57% have previously been identified in population surveys of human genetic variation (see [31] Table S4), along with 64% of L1 and 20% of SVA insertions, and most of the insertions were shared by at least two samples (Fig. S1B). We incorporated these germline insertions into sample-specific reference genomes using our TE-Seq pipeline[32]. This allowed us to map sequencing reads to non-reference elements and to prevent the misalignment of these reads to reference elements.

Next, we sought to identify putative somatic private insertion events. Unlike germline events, these events would be supported by only a single read (or, for a somatic event that occurred during development and was clonally amplified, very few reads). After stringent filtering (Fig. S1) to remove calls attributable either to read misalignments or non-TPRT derived somatic events, as well as manual inspection of insert sequence characteristics, we identified one putative somatic AluY insertion (in sample CTRL3) and one somatic L1HS insertion (in sample AD5) (Fig. 1B). Both insertions were supported by a single read, leaving open the possibility that these insertions occurred during development or adulthood. The AluY insertion is full-length and is located in an intergenic region (chr4: 159,407,307). The L1HS insertion is 5’ truncated and occurred in the second intron of the *NRXN3* gene (chr14: 72,851,739), which encodes a protein involved in synaptic cell-cell adhesion[33]. For each insertion, we show the insert sequence as well as a genome browser view of the read pileup at the insert site (Fig. 1C,D). Both insertions exhibit all of the hallmarks of TRPT-mediated retrotransposition. Each has a 3’ poly(A) sequence, as well as a target site duplication (TSD) of length consistent with TPRT. Furthermore, the putative cut-site sequence for these two events (GT/AAAT and AT/AAAG) closely resembles the known ORF2 cut-site motif preference TT/AAAA (Fig. S2C)[34, 35]. Sequencing read pileups at these events reveal two highly mappable loci and insert spanning reads (boxed in blue) that have typical error profiles (numbers of mismatches/small indels) for Nanopore DNA sequences. Additional plots documenting read phasing and local sequence homology for these events are provided in the supplement (Fig. S2).

An estimate of the expected number of events per cell can be computed by dividing the observed number of somatic events by the total number of diploid genomes sequenced. We obtained a value of 0.01, or 1 in 100 diploid genomes bearing a putative somatic insertion. As positive and negative controls, we applied our somatic insertion calling workflow to N2102Ep embryonal carcinoma cells, which express high levels of L1 ribonucleoprotein particles (RNPs)[36], and a human lung fibroblast cell line that does not express L1 elements[16]. We identified 10 insertions in N2102Ep cells at a coverage of 27.4x (0.73 events per diploid genome), and zero insertions in LF1 cells at a coverage of 164.7x (<0.006 events per diploid genome). N2102Ep insertions comprised four full-length, and five truncated, L1HS insertions, and one truncated Alu insertion (Fig. S3).

### Genome-wide examination of CpG methylation by Nanopore DNA-seq

Next, we sought to characterize changes to the CpG methylome in AD and examine RTE methylation in the aged brain. Mean global methylation varied among samples, ranging from 55% to 66% (Fig. 2A). We modeled CpG methylation using a binomial mixed-effects generalized linear model (GLM), with fixed effects for disease condition, age, and sex, and random intercepts for sample. Average methylation was significantly lower in AD (57.6%) than in CTRL (60.8%), (*p* = 0.048). In contrast, there was little variation in average methylation at gene promoter CpG islands (Fig. 2B), and the small difference accounted for by disease status was not significant (*p* = 0.74).

**Fig. 2.**
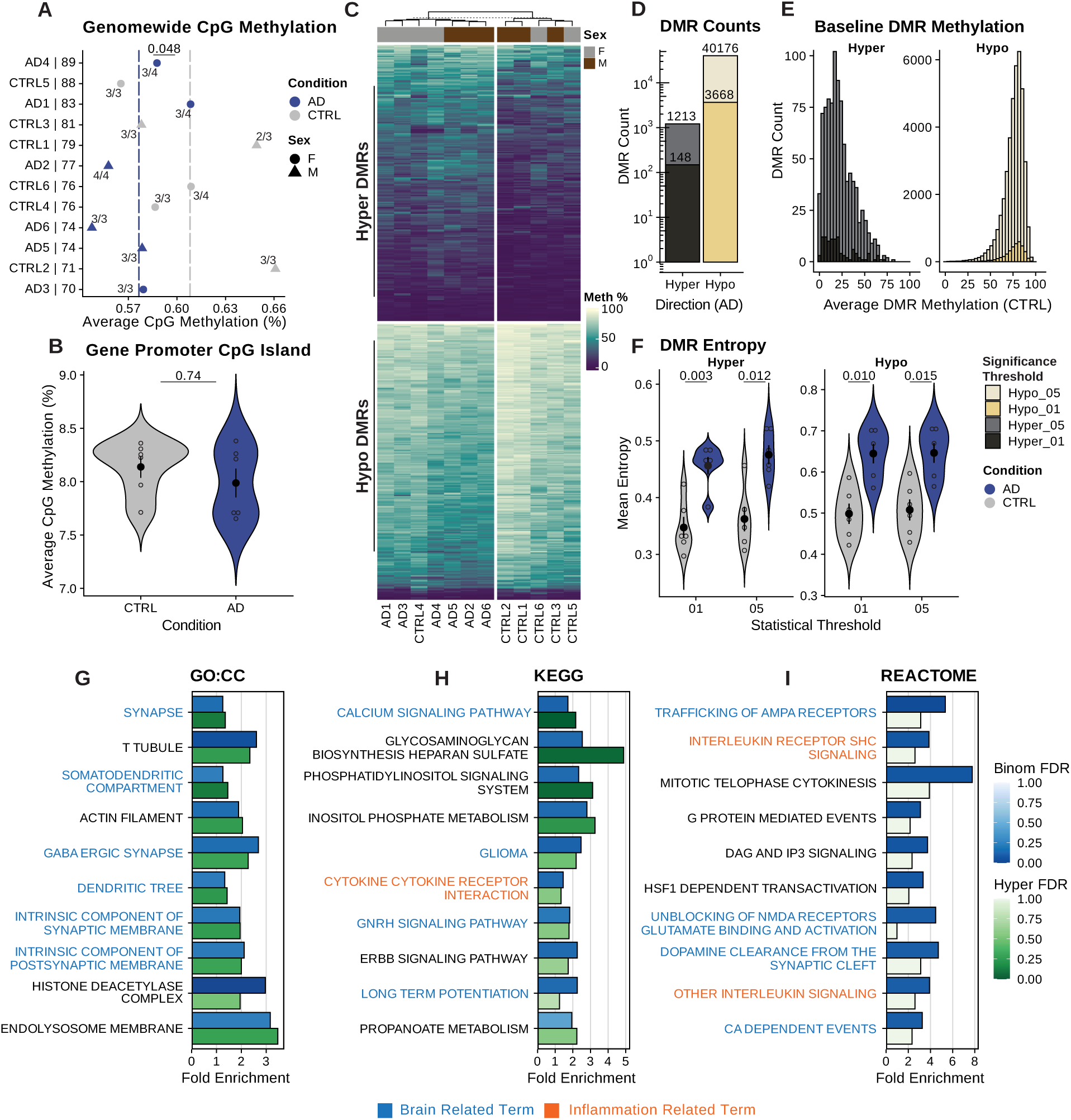
DNA CpG methylation differs between CTRL and AD samples. High coverage whole genome CpG 5-methylcytosine levels (5mCG) were determined by Nanopore DNA-Seq. (**A)** Scatter plot showing genome-wide, global mean methylation levels across all CpGs. Dashed vertical lines indicate the mean methylation level for a given disease condition. Samples are ordered by age (which is denoted after the underscore on the y-axis labels), colored by disease condition, and shaped according to sex, and *APOE* genotype is shown (e.g. AD4 – *APOE* 3/4). Statistical analysis was performed using a binomial mixed-effects model with fixed effects for disease condition, age, and sex, and random intercepts for sample. A significant effect of condition was observed (*p* = 0.048). (**B**) Violin plot showing the distribution of average CpG methylation in gene promoter (−5 kb to +1kb relative to TSS) CpG islands. Statistical analysis was performed using a binomial mixed-effects model with fixed effects for disease condition, age, and sex, and random intercepts for sample and gene. Disease condition did not have a significant effect on methylation (*p* = 0.74). (**C**) Heatmap of the mean CpG methylation for the top 500 hyper/hypo methylated DMRs, as ranked by p-value. Rows and columns were hierarchically clustered. (**D**) Bar plot indicating the number of differentially methylated regions (DMRs) grouped by CpG island status and colored by direction of methylation change in AD relative to CTRL. DMRs were called using the DSS statistical program wherein methylation was modeled as a function of disease condition, adjusting for age and sex. Before merging adjacent DMLs into DMRs, DMLs were filtered with two p-value thresholds, 0.05 and 0.01. (**E**) Histogram showing the mean CTRL methylation of all DMRs, faceted by direction of change relative to CTRL. (**F**) Violin plot showing the distribution of DMR methylation entropy. Samples are colored by disease condition. (**G-I**) Functional enrichment of promoter/enhancer overlapping DMRs was assessed using the rGREAT statistical program, using all promoter/enhancer regions as background. mSigDB gene set subcollections were queried for enrichment. We show the top 10 enriched sets for the GO: Cellular Component, KEGG, and REACTOME subcollections, as ranked by the mean of their FDR-adjusted binomial/hypergeometric p-values.

To investigate the presence of differentially methylated regions (DMRs) in AD, we used the DSS statistical program, which uses a Bayesian GLM to model methylation. DSS first calls differentially methylated loci (DMLs) and then merges DMLs passing a statistical significance threshold (here we used two thresholds, *p* ≤ 0.05 and *p* ≤ 0.01) into DMRs. Adjusting for age and sex, disease condition was associated with 41,389 DMRs at the 0.05 threshold, and 3,816 DMRs at the 0.01 threshold (Fig. 2D) (see Methods for statistical thresholds). Concordant with the global decrease in methylation observed in AD samples, these regions were predominantly hypomethylated. At the 0.05(0.01) p-value threshold, these regions averaged 399(317) bp in length and encompassed 7.1(7.2) CpG sites.

We plotted a heatmap of the 500 most differentially methylated hyper/hypo DMRs, split samples into two groups using k-means, and clustered rows and samples by hierarchical clustering (Fig. 2C). As expected, samples largely clustered by biological condition. The absolute value of the percent change in methylation across hypomethylated DMRs was 11.3(14.7)% and, for hypermethylated DMRs, 10.4(16.4)%. DMRs hypermethylated in AD tended to be poorly methylated in CTRL samples, whereas DMRs hypomethylated in AD tended to be robustly methylated in CTRLs (Fig. 2E). This suggests a process of epigenetic drift, in which low-entropy regions (CpGs are either mostly methylated or demethylated) become more disordered with time, netting out a directional shift in methylation. Indeed, average methylation entropy was increased in both hypo- and hyper-DMRs in AD (Fig. 2F; *p* < 0.05 for all contrasts). Statistical significance was assessed with a linear model regressing per-sample mean DMR entropy on disease condition while controlling for age, sex, and mean read depth per DMR (see Methods for details of entropy calculation).

We queried the DMRs for overlap with CpG island and chromatin state annotations: CpG islands, shores (2 kb regions flanking islands), shelves (2 kb regions flanking shores), and “open sea” regions (Fig. S4A). Most DMRs did not occur in or adjacent to CpG islands, though they were strongly enriched in islands when considering the fraction of the genome occupied by these regions. DMRs were predominantly found in B compartment-associated chromatin states, especially in quiescent, Polycomb-repressed, and heterochromatic regions, and these DMRs tended to be hypomethylated (Fig. S4B). This was largely due to the vast size of these regions; hypo-DMRs were only weakly enriched in these regions when accounting for their genomic footprint. ZNF/Repeat regions, which are enriched for RTEs, were two orders of magnitude more likely to overlap hypo- versus hyper-DMRs. In contrast, transcription start site (TSS) associated states overlapped comparable numbers of hypo- and hyper-DMRs and were enriched for hyper-DMRs and depleted of hypo-DMRs.

We next examined gene sets for association with DMRs occurring in promoters (the −5 kb to +1 kb region surrounding a gene TSS) or enhancers. Overall, promoters and enhancers primarily overlapped hypo-DMRs, but enriched for hyper-DMRs (Fig. S4C). We used rGREAT[37] to query the Molecular Signatures Database (mSigDB) gene set subcollections for association with our DMRs. We show enrichment results for the top 10 most enriched GO Cellular Component, KEGG, and REACTOME terms for DMRs called using the 0.05 threshold (Fig. 2G-I). rGREAT performs two types of enrichment tests, one based on a binomial distribution and the other based on a hypergeometric distribution. We ranked gene sets according to the mean of their binomial and hypergeometric enrichment FDR-adjusted p-values. Most of our enriched terms could be classified as either brain-related or inflammation-related. “Synapse,” “Calcium Signaling Pathway,” and “Trafficking of AMPA Receptors” were the top enriched terms in these three subcollections.

In sum, global genome methylation experiences a hypomethylation trend in AD. Differentially methylated regions see increased methylation entropy in AD, suggesting increased epigenetic drift in AD. Repressive genomic compartments see the greatest degree of hypomethylation, whereas hyper-DMRs are enriched in gene cis-regulatory elements. Gene sets important to brain function and inflammation are most enriched by our differentially methylated regions.

### Young L1s, and RTEs more broadly, are hypomethylated in AD

RTEs were not exempt from overlap with hypo-DMRs. We queried our DMRs for overlap with the promoter regions of evolutionarily young, full-length (FL, >95% of consensus length) RTEs. We examined the 5’ untranslated region (UTR) of L1 elements (L1HS-L1PA6), the 5’ long terminal repeat (LTR) region flanking human ERV elements (HERVK and HERVL), and the entire ∼300 bp Alu element (AluY). L1 subfamilies had the highest fraction of their members overlap hypo-DMRs (Fig. 3A). Alu and L1 subfamilies were enriched for hypo-DMRs and depleted of hyper-DMRs (though given the low number of hyper-DMRs, and the limited genome fraction occupied by these young RTE subfamilies, the extent of depletion is difficult to estimate), whereas the relationship of ERVs to DMRs was less clearly defined (Fig. 3B). We next plotted the average promoter methylation for every full-length member of these subfamilies in each of our samples (Fig. 3C). We observed a reduced median promoter methylation across these young RTE subfamilies. In both AD and CTRL, a subset of elements stands out as having particularly low average promoter methylation.

**Fig. 3.**
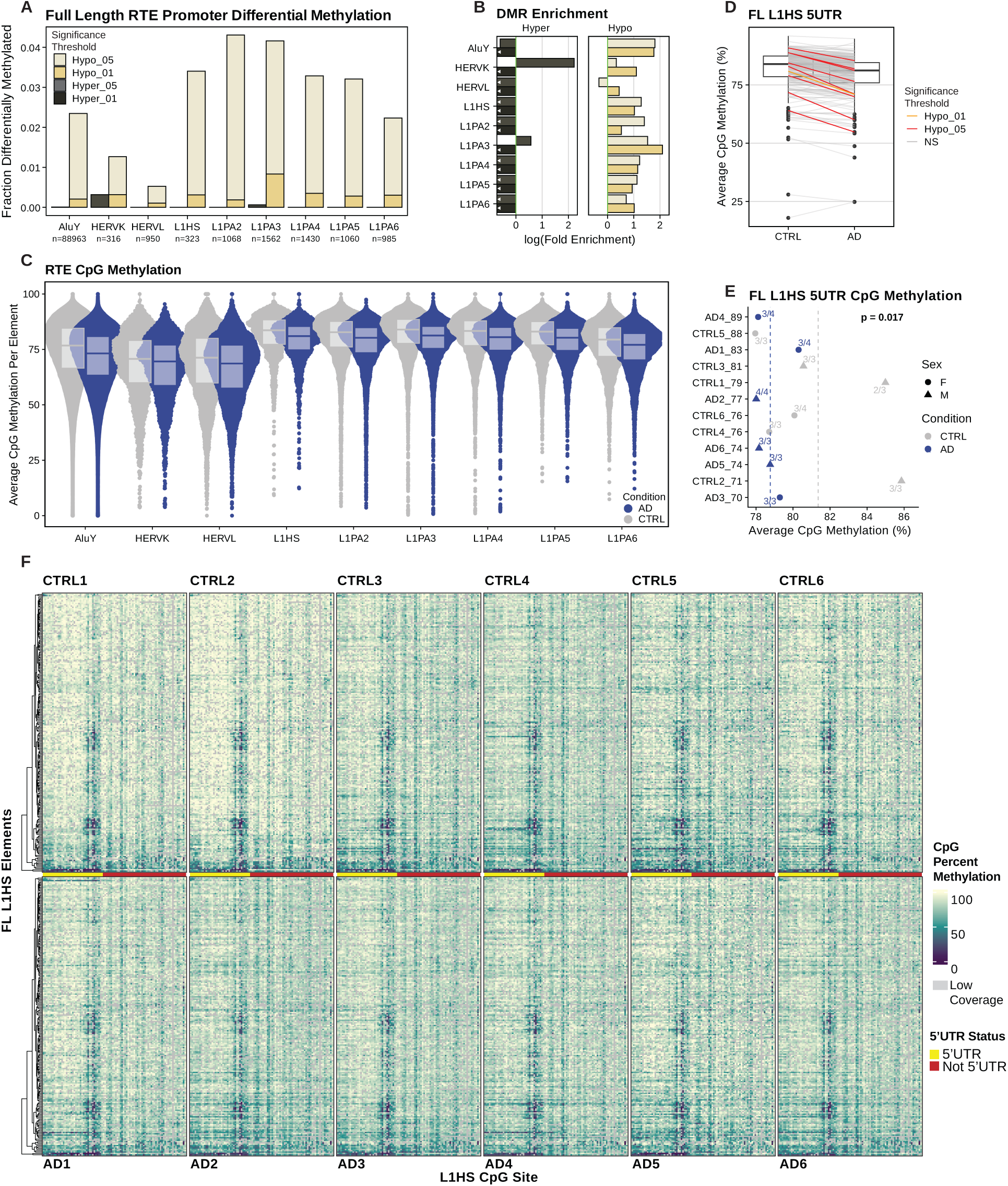
Young RTEs are hypomethylated in AD. (**A**) Bar plot showing the fraction of RTE subfamily members whose transcriptional control region overlaps a DMR. For AluY and LTRs, the transcriptional control region is the element itself; for L1 subfamilies, this is the 5’ UTR or first 909 bp, colored by direction of change relative to CTRL. Only FL elements (>95% of consensus element length) are considered. (**B**) Bar plot of the natural logarithm of the fold enrichment of DMRs in RTE subfamily transcriptional control regions. Bars with a white arrow pointing left and touching the y-axis go to negative infinity. We modeled the expected fraction of DMRs occurring in a set of regions as a binomial with probability equal to the genome fraction occupied by these regions. (**C**) Strip plot showing average methylation levels within individual FL RTE element transcription control regions, grouped by RTE subfamily and colored by disease condition. (**D**) Paired scatter plot showing changes in 5’ UTR methylation levels for all FL L1HS elements between disease conditions. Line segment color indicates whether the 5’ UTR overlaps a DMR as well as the direction of methylation change. (**E**) Scatter plot showing mean FL L1HS 5’ UTR methylation levels. Dashed vertical lines indicate the mean methylation level for a given disease condition. Samples are ordered by age (which is denoted after the underscore on the y-axis labels), colored by disease condition, and shaped according to sex, and *APOE* genotype is shown (e.g. AD4 – *APOE* 3/4). Statistical analysis was performed using a binomial mixed-effects model with fixed effects for disease condition, age, and sex, and random intercepts for sample and L1HS locus. A significant effect of condition was observed (*p* = 0.017). (**F**) Heatmaps showing the mean CpG methylation for all reference FL L1HS elements (top: CTRL samples, bottom: AD samples). Heatmap rows represent individual L1HS elements and are hierarchically clustered. Heatmap columns represent individual CpG sites and are ordered by sequence position. Row and column order are preserved among all heatmaps.

For FL L1HS elements, the changes in aggregate methylation in AD are most driven by a substantial minority of elements that overlap hypo-DMRs (Fig. 3D). We fit a binomial mixed-effects model to test for differences in CpG methylation at the FL L1HS 5′ UTR. CpG methylation was modeled using fixed effects for disease condition, age, and sex, and random intercepts for sample, L1HS locus, and CpG ID (e.g. CpG_1 for the first CpG in the L1HS 5’ UTR). CpG methylation was significantly decreased in AD compared to CTRL, *p* = 0.017 (Fig. 3E). A post hoc model including an interaction between disease condition and sex revealed that the relationship between methylation and AD is modulated by sex. Specifically, control males appeared most protected against reductions in FL L1HS methylation (*p* < 0.001). A heatmap showing CpG methylation by CpG site for all reference (Fig. 3F) and non-reference (Fig. S5) FL L1HS elements revealed striking similarities among samples, and substantial variation among L1HS loci. Roughly 40% of all reference elements (which are hierarchically clustered and whose ordering is preserved between heatmaps) have a pronounced region of lower methylation in the 5’ UTR (found at position 363 bp – 767 bp), which roughly corresponds to the L1 antisense promoter (400 bp – 600 bp). Given that this region of lowered methylation is shared among all samples and across so many L1HS elements, we suspect that this reduced methylation is not sufficient to trigger L1 sense strand derepression and ORF1/ORF2 expression. Conversely, the first 328 bp of most elements were heavily methylated, leading us to suspect that expression may be most sensitive to CpG methylation in this region. We found it noteworthy therefore that a minority of elements found at the bottom ∼5% of each reference element heatmap (Fig. 3F) had low average methylation values (<60%) across the entire 5’ UTR region, including in this 328 bp region.

Two full-length elements (L1HS_2p13.2_1 and L1HS_2q21.1_2) stand out as being highly demethylated across all samples. L1HS_2p13.2_1 is located in a *ZNF638* intron, in an antisense orientation, and encodes intact ORF1 protein and a partial ORF2 open reading frame. This 3’ truncated ORF2 reading frame spans 1,518 bp and includes the entire ORF2 endonuclease (EN) domain before reaching a premature stop codon. L1HS_2q21.1_2 is intergenic, and retains only a partial, 5’ truncated ORF2 open reading frame, which spans 3,033 bp and includes the full reverse transcriptase (RT) domain. We examined ENCODE ChIP-seq tracks at these elements focusing on H3K9me3, H3K27me3, and H3K27ac. L1HS_2p13.2_1 lacked enrichment of the repressive histone marks H3K9me3 and H3K27me3 and was enriched for H3K27ac specifically at the element’s 5’ UTR (including for the only brain-resident primary cell type profiled in this ENCODE dataset, brain microvascular endothelial cells, or BMECs) (Fig. S6C). In fact, L1HS_2p13.2_1 was the only FL L1HS (of a total of 332 FL elements) whose 5’ UTR overlapped an H3K27ac peak in BMECs, in stark contrast to L1HS_2q21.1_2 which was enriched for repressive histone marks and depleted of H3K27ac at its 5’ UTR. The consistent loss of CpG methylation observed in our Nanopore data, along with the enrichment of active histone marks in ENCODE data, suggests L1HS_2p13.2_1 may be a particularly transcriptionally active L1HS element and warrants enhanced scrutiny. Concordantly, our RNA-Seq analysis (see Methods for more details) revealed that L1HS_2p13.2_1 (Fig. S7A) was one of the most highly expressed FL L1HS elements across samples (94^th^ expression percentile), whereas L1HS_2q21.1_2 (Fig. S7B) was poorly expressed (40^th^ expression percentile).

### Highly demethylated reads increase in frequency in AD at L1HS CpG promoters

We next sought to disaggregate bulk CpG methylation into individual reads overlapping full-length L1HS elements. This read-centric approach is valuable in that it can reveal changes in CpG methylation at a given locus in individual cells. While a partial loss of CpG methylation at an L1HS locus may be attributable to increasing epigenetic drift in a highly methylated region, total loss of CpG methylation at a locus in an individual cell is far likelier to have functional consequences. Across FL L1HS promoters, the majority of reads were highly methylated (>75% methylation). Notably, we also observed elements in which individual reads were nearly fully demethylated (Fig. 4F). To thoroughly characterize changes in the fraction of highly demethylated reads (HDRs), we focused on three regions (Fig. 4A) that we reasoned would be important for the transcriptional regulation of L1HS elements: the 5’ UTR (first 909 bp), the CpG island within the 5’ UTR (first 500 bp), and the region of consistently elevated methylation within the CpG island we identified (Fig. 3F; first 328 bp). Across these regions, we binned reads into methylation quartiles and considered reads less than 50% methylated to be highly demethylated. For all regions, the overall fraction of HDRs increased in AD in at least one of the two HDR quartiles (Fig. 4B). Statistical analysis was performed using binomial mixed-effects models with fixed effects for disease condition, age, and sex, and random intercepts for sample and L1HS locus. An effect was considered significant at FDR-adjusted *p* ≤ 0.05.

**Fig. 4.**
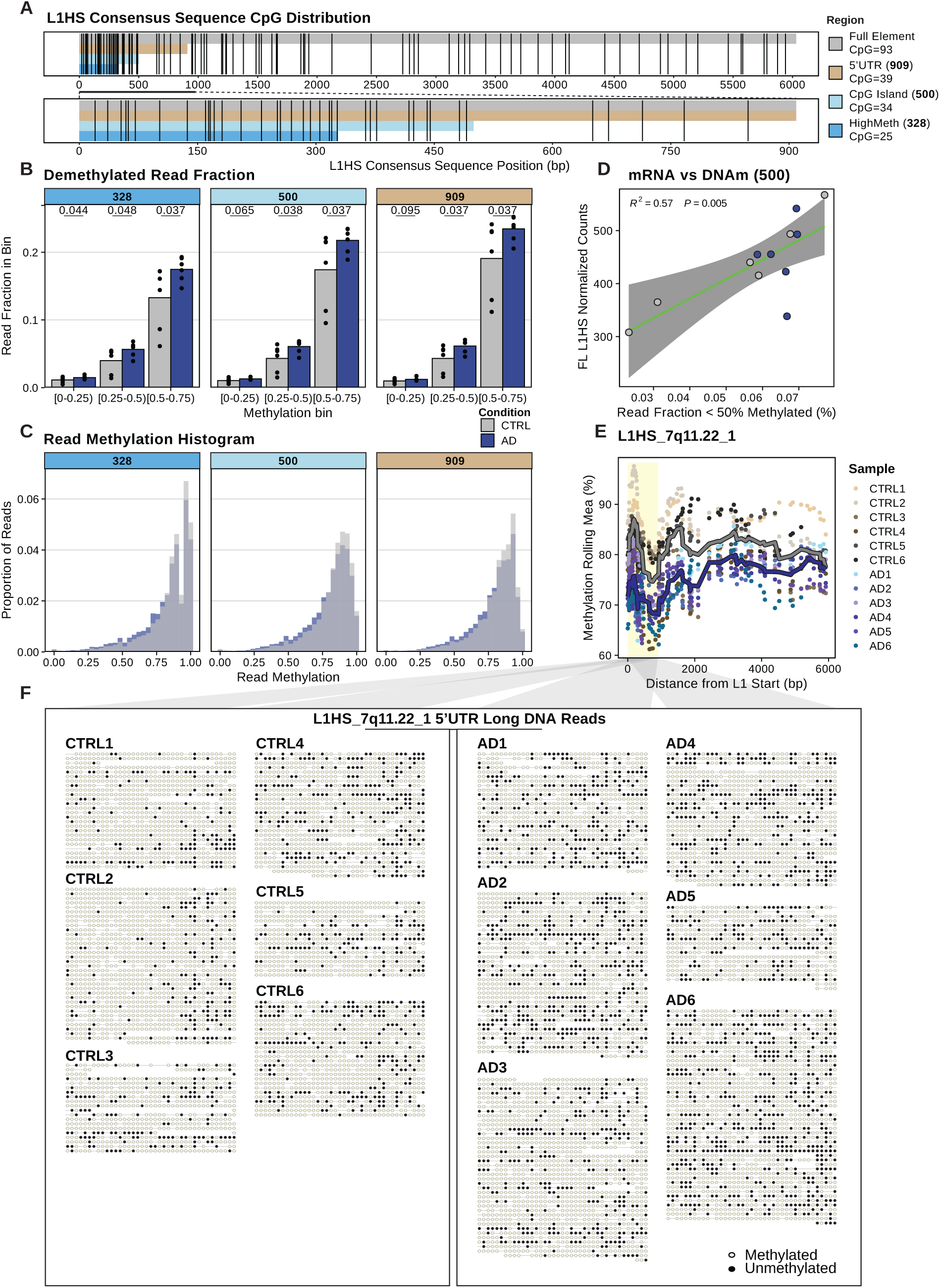
FL L1HS 5’ UTR hypomethylation is mediated by a sharp loss of methylation in a subset of cells. (**A**) Distribution of CpG sites in the consensus L1HS element. Shaded regions denote the full element in gray, the internal promoter 5’ UTR in brown (first 909 bp), the CpG island within the 5’ UTR in light blue (first 500 bp), and the region of consistently high methylation identified in Fig. 3C in blue (first 328 bp). (**B**) Bar plots showing the fraction of reads overlapping the first 328, 500, or 909 bp of the FL L1HS 5’ UTR binned by their average CpG methylation. For inclusion, reads must cover at least 75% of the CpGs found in the region. The effect of disease condition on the fraction of reads in each bin was estimated using a binomial mixed-effects model, with fixed effects for condition, age, and sex, and random intercepts for sample and L1HS locus. We report FDR-adjusted p-values. (**C**) Histogram showing the distribution of read methylation over the first 328, 500, or 909 bp of the FL L1HS 5’ UTR. (**D**) We performed stranded, paired-end RNA-seq on rRNA-depleted RNA extracted from the same brain region used for Nanopore DNA-seq in each of our samples. FL L1HS expression is plotted against the highly demethylated read (<50% methylated) fraction over the first 500 bp of the 5’ UTR. Statistical analysis was performed using a linear model regressing FL L1HS expression against HDR fraction (R^2^ = 0.57, *p* = 0.005). (**E**) Scatter plot showing the average methylation level by CpG for all CpGs in the L1HS_7q11.22_1 element. The 5’ UTR region is shaded in yellow. Condition averages are shown as lines. (**F**) Bubble plot showing all reads overlapping the L1HS_7q11.22_1 element’s 5’ UTR (region shaded in yellow). Reads are trimmed to not extend beyond this region. Each line segment represents a single sequencing read and each bubble represents a CpG site. The CpG is methylated if yellow, and demethylated if black.

The distribution of read-level methylation (Fig. 4C) was left-skewed. To quantify the degree to which methylation probability varied across reads (suggesting cell-of-origin effects beyond sampling variation) we fit a beta-binomial model in which the precision parameter (φ) describes how tightly methylation probabilities cluster around the mean. φ was modeled as a function of disease status and random effects were included for sample, L1HS locus, their interaction, and CpG ID (e.g. CpG_1 for the first CpG in the L1HS 5’ UTR). For all 5’ UTR regions, we observed significantly lower precision (higher overdispersion) in AD (likelihood ratio test, *p ≤* 0.0001). Higher φ indicates less heterogeneity among reads (closer to a binomial model), whereas lower φ reflects greater heterogeneity (overdispersion). We estimated φ at 13.61 (95% CI: 13.57–13.66) in controls and 9.68 (95% CI: 9.63–9.74) in AD samples, indicating greater read-level variability in AD. We visualize the effect of these values of φ on the distribution of read methylation for a hypothetical locus with methylation probability equal to 0.75 in both CTRL and AD (Fig. S8D). These values of φ are consistent with additional cell-of-origin effects in AD. One possibility is that a subset of cells exhibits loss of methylation across numerous L1HS loci. Alternatively, a larger number of cells may lose methylation at a smaller number of loci. Both scenarios are consistent with the observed methylation overdispersion, especially considering the increase in HDRs observed at L1HS 5’ UTRs in AD.

We next looked for methylation changes at individual FL L1HS loci. Only a small number of L1HS loci showed significant changes in the fraction of HDRs using a nominal p-value threshold of 0.05 (no loci were significant after FDR correction) (Fig. S8B). This was largely due to high inter-sample variation in the fraction of HDRs within conditions. At a given locus, often only one or two samples exhibited an appreciably increased number of HDRs. Most of the gains in HDRs in AD occurred at highly methylated loci, and AD samples had fewer loci with no HDRs (Fig. S8A). The L1HS_7q11.22_1 5’ UTR (Fig. 4E) recapitulates the broader patterns noted above. Several AD samples exhibited a substantial loss of methylation at this element’s 5’ UTR, and a read-centric view reveals that this decrease is primarily driven by an increase in the number of HDRs in samples AD2, AD4, and AD6 (Fig. 4F). Furthermore, variability in L1HS_7q11.22_1 read methylation is greater in AD than CTRL (Fig. S8E).

Highlighting the sensitivity of our approach, we resolved allelic effects on methylation at a polymorphic L1HS locus in sample AD1 (Fig. S10A). We identified a heterozygous 5 kb deletion in the body of the L1HS_1q31.3_4 element that left its 5’ UTR intact. Those reads bearing the 5 kb deletion showed substantial demethylation in the 5’ UTR whereas reads lacking this deletion showed a robustly methylated 5’ UTR (Fig. S10C). This finding suggests that primary sequence motifs within this L1HS promoter are insufficient to initiate or maintain DNA methylation in the absence of a downstream, non-intronic, long transcriptional unit. This is consistent with the known role of the HUSH complex, which silences long (>1 kb) intronless transcripts, such as young, full-length L1 elements[38]. That this 3’ truncated element would evade such targeting is therefore highly plausible and lends credence to our broader approach. Of note, L1HS_1q31.3_4 is located in the 3^rd^ intron of the *DENND1B* gene which has recently been identified in a canine and human genome-wide association study (GWAS) as having intronic SNPs associated with obesity[39]. Our methylation data suggest that the 5 kb deletion promotes 5’ UTR demethylation, which we speculate may affect *DENND1B* expression via L1HS 5’ UTR cis-regulatory activity[40]. That this variant was identified in one of only 12 individuals sequenced suggests it may be present at an appreciable allelic frequency in the general human population.

### DNA methylation predicts gene expression

To assess whether the observed changes in promoter methylation were associated with gene expression, we performed stranded 150 bp paired-end RNA sequencing on samples from the same donors and brain region used for Nanopore DNA sequencing (Fig. S9). The top 30 terms identified in our enrichment analysis of DMRs were more likely to be enriched in our RNA-seq GSEA than were other terms. Of these 30 gene sets, 14 (47%) were enriched at the transcriptional level based on an FDR *≤* 0.1 cutoff. This overlap corresponds to a fold enrichment of 1.61 (hypergeometric test, *p* = 0.029). Of these doubly enriched gene sets, the top RNA-enriched term in each gene set ontology was “GO:CC Synapse”, “KEGG Cytokine Cytokine Receptor Interaction”, and “REACTOME Trafficking of AMPA Receptors”. Gene set enrichment analysis of the mSigDB Hallmark collection (Fig. S9B) identified a strong inflammatory signature in our AD sample transcriptome. The top six terms enriched in AD were inflammation-related gene sets and included the “Interferon Alpha Response” gene set. We have previously shown that L1HS activity induces a type I IFN response[16].

FL L1HS elements were more highly expressed in AD than in control samples, though this difference did not reach statistical significance (negative binomial GLM with fixed effects for disease condition, age, sex, RIN; DESeq2-derived size factors were provided as an offset, *p* = 0.644) (Fig. S9E). FL L1HS expression was significantly associated with the fraction of highly demethylated reads overlapping any of the three 5’ UTR regions (Fig. 4D - 5’UTR CpG island, first 500 bp: R^2^ = 0.57, *p* = 0.005), as well as with average CpG methylation in these regions (Fig. 4D, S8C). This finding suggests that the changes in methylation we observed across samples are responsible for differences in FL L1HS expression.

Overall, we find good concordance between promoter methylation changes and transcriptional signatures for both RTEs and genes. These findings suggest that long read methylation profiling can provide meaningful insight into transcriptional changes in aged brain samples and may be particularly advantageous in other contexts where RNA integrity or availability is limiting.

## Discussion

After highly stringent filtering (Fig. S1), we report putative somatic TPRT-mediated insertions in the aged human brain using Nanopore DNA-sequencing (Fig. 1D-H; Fig. S2; Fig. S3). These events capture all hallmarks of retrotransposition: a poly-A tail, target site duplication, and ORF2 cut site motif and are either full-length or 5’ truncated. Across all samples, we sequenced a total of 204.6 diploid genomes and identified two high-confidence somatic insertions; therefore, we estimate the per-cell prevalence of somatic TPRT-mediated insertions to be of ∼0.01. In contrast to earlier reports of ubiquitous somatic mosaicism in the human brain[41], our estimate is consistent with a body of literature identifying low rates of somatic L1 insertions in the human brain. Previous work derived from single cell short-read sequencing of MDA amplicons estimated somatic L1 insertion prevalence to range from 0.04 – 0.29 insertions per cell in the human brain[42–44]. Our putative somatic insertion events were supported by a single sequencing read. While this is compatible with these insertions having occurred either during or post-development, our examination of CpG methylation at L1HS promoters suggests that these insertions may well have occurred in advanced age.

Global CpG methylation was decreased in AD (Fig. 2A). Methylation entropy was increased at DMRs, suggesting a process of epigenetic drift in AD (Fig. 2E,F). While aggregate CpG methylation was unchanged across gene promoter CpG islands (Fig. 2B), we identified thousands of differentially methylated regions overlapping gene promoters and enhancers (Fig. S4B). Inflammatory and brain function-related gene sets were most associated with DMRs (Fig. 2G-I). The most enriched gene sets included “Synapse,” “Calcium Signaling Pathway,” and “Trafficking of AMPA Receptors.”

On a per-element basis, the transcriptional control region of evolutionarily young, full-length L1 elements most frequently overlapped hypomethylated DMRs compared to other young RTE subfamilies (Fig. 3A). Given the known importance of CpG methylation in suppressing young L1HS expression[45, 46], we found it striking that 5% of full-length L1HS promoters were poorly methylated (<60% average CpG methylation) across both control and AD samples, that 7% of reads overlapping the L1HS promoter were less than 50% methylated, and that 1% of reads were less than 25% methylated (Fig. 4B). The frequency of highly demethylated reads (HDRs) at FL L1HS promoters was increased in AD (Fig. 4B). These gains in HDRs tended to come from highly methylated loci at which no highly demethylated reads were observed in CTRL samples (Fig. S8A), suggesting that maintenance of methylation may be particularly important at these loci. FL L1HS elements with increased HDRs tended not to be shared among AD samples and were instead unique to one or several samples, painting a stochastic picture of L1HS derepression (Fig. S8B). Consonantly, read methylation at FL L1HS promoters was overdispersed, and the degree of overdispersion was higher in AD. In other words, the frequency of highly demethylated reads at a given locus in a given individual is higher than would be expected from a simple Bernoulli process (where read methylation is modelled as a series of coin flips in which the methylation probability for each CpG is fully determined, i.e. constrained, by the sample, L1HS locus and CpG ID), suggesting that there are cell-or-origin effects on L1HS 5’ UTR methylation. This is consistent with certain rare cell types or cell states (e.g. cellular senescence) disproportionately comprising the source of the highly demethylated reads sequenced. CTRL men were the most protected against decreases in L1HS methylation (Fig. 3E). Alongside the notion of L1 involvement in AD pathogenesis, this finding is consistent with greater female risk for developing the disease[47]. Overall, we find that repression by CpG methylation is compromised in a subset of FL L1HS elements in the aged brain and that this effect is exacerbated in AD.

Supporting the importance and biological relevance of CpG methylation in controlling L1HS expression, we observed a strong negative relationship between aggregate L1HS promoter methylation and L1HS expression (Fig. 4D, S8C). While Lanciano et al.[48] also observed this association, they showed that CpG demethylation at an L1 promoter does not guarantee expression. It is likely that those L1s that are demethylated but remain unexpressed reside in regions marked by repressive histone modifications. We found two L1HS elements, L1HS_2p13.2_1 and L1HS_2q21.1_2, that were particularly demethylated across all samples.

We found it quite notable, therefore, that L1HS_2p13.2_1 was also found in a transcriptionally permissive histone environment in ENCODE data (Fig. S6C). Indeed, L1HS_2p13.2_1 was the only FL L1HS element whose 5’ UTR harbored an H3K27ac peak in BMEC (primary brain micro epithelial) cells, the ENCODE-profiled cell type most pertinent to our present work.

Consistently, we identified L1HS_2p13.2_1 as one of the most highly expressed L1HS elements (94^th^ percentile among FL elements) in our RNA sequencing analysis (Fig. S7A). L1HS_2p13.2 has previously been identified as a transcriptionally active element in adult human neurons[49] and has been shown to be poorly methylated in several tissues[31]. L1HS_2p13.2_1 encodes an intact ORF1 and a 3’ truncated ORF2 lacking the reverse transcriptase (RT) domain. This 3’ truncated ORF2 nevertheless possesses an intact endonuclease (EN) domain, leaving open the possibility that it could promote genomic dsDNA breaks in the aged brain. In fact, a recent report suggests that 3’ truncation of ORF2 can enhance endonuclease activity towards genomic DNA[50]. DNA damage, in particular double stranded breaks, have been linked to neurodegeneration[51–53].

That the L1HS_2p13.2_1 element resides in the ZNF638 gene, which binds to retroelement DNA sequences and silences them through recruitment of the HUSH complex[54–56], invites further scrutiny. It has been shown that L1HS_2p13.2_1, through its anti-sense promoter (ASP), can generate a chimeric ZNF638 transcript [49, 57]. We note that the L1-ZNF638 isoform comprises the ZnF and DNA-binding domains, but lacks the N-terminal 471 amino acid region which was shown to be sufficient for its association with the HUSH complex (specifically, for its affinity to MPP8)[56]. We therefore speculate that expression of this L1-ZNF638 isoform may promote RTE derepression by (i) failing to recruit HUSH to target loci, and (ii) competing with HUSH-competent, full-length ZNF638 for RTE binding. The hypomethylation of the L1HS_2p13.2_1 ASP we observed in the aged brain may therefore encourage RTE derepression via this hypothesized mechanism.

In sample AD1 we identified a heterozygous, 5 kilobase deletion in the body of a FL L1HS element which left the 5’ UTR intact. Sequencing reads lacking the deletion show a highly methylated 5’ UTR, whereas those with the deletion show a highly demethylated 5’ UTR (Fig. S10C). This finding suggests that HUSH complex action, which depends on long, intronless transcriptional units such as the body of an L1HS element, may be responsible for repression of this locus. This finding also highlights the ability of Nanopore long-read sequencing to resolve allelic effects on methylation at repetitive loci that would be challenging to examine with short-read sequencing approaches.

### Conclusions

In sum, somatic retrotransposition and DNA methylation provide two lines of evidence that suggest that L1HS elements are mobilized in the aged and AD brain. The identification of somatic Alu and L1 insertions provides strong evidence of LINE-1 ORF2-activity in the human brain. We report substantial numbers of highly demethylated reads at FL L1HS promoters and found this hypomethylation to be exacerbated in AD. Our findings are consistent with non-mutually exclusive models wherein (i) a rare class of cells sees a profound loss of methylation across many L1HS promoters, or (ii) a broad-based increase in epigenetic disorder is translated into many cells stochastically losing control over a limited number of L1HS elements. These scenarios support the notion that young L1 elements may challenge cellular homeostasis via genotoxic or inflammatory mechanisms in the context of AD[16].

## Methods

### Sample selection

De-identified frozen prefrontal cortex (PFC, Brodmann area 9) brain tissue from the gray matter of six AD cases was obtained from the Mount Sinai Brain Bank (NIH NeuroBioBank). The selected cases were clinically confirmed AD patients with dementia and post-mortem determination of Braak neurofibrillary tangle (NFT) stage V-VI, indicating neocortical involvement. Additionally, age- and sex-matched frozen PFC tissues (Brodmann area 9) from six cognitively normal individuals were used as controls. Sample CTRL5 could not definitively be assigned a Braak stage as no p-tau, b-amyloid, a-synuclein, or Bielschowsky stains were performed on any region. It was categorized as a control because of the lack of clinical indications for neurodegeneration and because no plaques, tangles, or Lewy bodies were identified on H&E-stained sections. All tissues were stored at −80°C until further processing.

### DNA extraction, library preparation, and sequencing

We isolated genomic DNA using the NEB Monarch Spin gDNA Extraction Kit. Sequencing libraries were prepared following the unmodified Nanopore DNA Ligation Sequencing Kit V14 (SQK-LSK114) protocol for a Promethion flow cell. Each sample was loaded onto its own 10.4.1 PromethION Flow Cell. Sequencing was performed on a P2-solo instrument, which allows for two flow cells to be run concurrently. This allowed us to balance sequencing runs with respect to disease condition (an AD sample would be run alongside a CTRL sample). Two sequencing libraries were prepared for each sample. Flow cells were washed using the flow cell wash kit (EXP-WSH004-XL) midway through sequencing and a fresh sequencing library derived from the same sample was loaded. Sequencing was controlled using the MinKNOW software version 24.02.6 and was allowed to run for three days or until nearly all sequencing pores were inactive.

### Nanopore basecalling and quality control

Nanopore sequencing data were basecalled using Dorado (VN:0.5.1+a7fb3e3) with the high-accuracy, 5mCG and 5hmCG basecalling model (dna_r10.4.1 e8.2 400bps hac@v4.3.0 5mCG_5hmCG@v1). Basecalled reads were then aligned to the telomere-to-telomere T2T-HS1 (GCF_009914755.1) human genome[58]. Quality control metrics were created using PycoQC[59]. A filtered BAM to be used in various sensitive downstream applications that require only high-confidence alignments was then created. We retained only those alignments with a read quality score ≥ 10, a MAPQ ≥ 10, and filtered out secondary and supplementary alignments using samtools[60] (samtools view -b -F 2304 -q 10 -e ‘[qs] >= 10’).

### Differentially methylated locus/region (DML/R) calling

BedMethyl files were generated for all samples from the filtered BAMs using the Oxford Nanopore Technologies (ONT) modkit pileup command. CpG methylation data were then fed to the DSS[61] program in R for differential methylation analysis. DSS uses a Bayesian GLM regression framework to infer differentially methylated loci (DMLs). We modeled methylation as a function of disease condition and included sex and age as covariates. DSS subsequently merged adjacent statistically significant DMLs to obtain differentially methylated regions (DMRs). Since DMR calling is sensitive to the p-value threshold used to filter DMLs, DSS recommends users try multiple thresholds. Accordingly, we report all DMR-based results for two p-value thresholds, 0.05 and 0.01.

### Methylation analysis

Our methylation analyses took on three broad flavors: DMR based, CpG position based, and read-based. In all cases, we excluded from our analysis sequences on the mitochondrial and the Y chromosome.

DMR based analyses overlap DSS called DMRs with genome annotations in R using the Genomic Ranges package[62]. An 18-state chromHMM annotation of the frontal cortex (derived from two females aged 67 and 80 years) was obtained from the ENCODE data portal[63] (ENCFF927GIF). Coordinates were lifted over from Hg38 to T2T-HS1 using the UCSC liftOver tool[64]. To quantify methylation entropy at DMRs, we used the ONT modkit entropy command, which computes methylation entropy (ME) as:

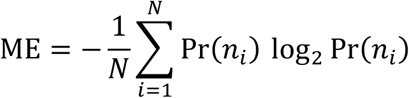

where N is the set of all methylation patterns and Pr(n_i_) is the empirical probability of that pattern.

CpG-based analyses average methylation levels across all reads covering a CpG. Subsequently, methylation levels across CpGs in each region or set of regions can be modeled across disease condition using a binomial mixed-effects models, adjusting for sex and age. It is critical that samples share the same set of CpGs and that these sites have sufficient coverage for a reliable methylation estimate. We filtered CpG sites to only consider those that have a coverage of 7 or more in each sample, resulting in 24.3 million sites included in our analysis (out of 32.3 million total sites in the genome).

Read-based analyses began by collecting all reads that overlapped a given region, or set of regions, of interest (ROI). Having observed that a small fraction of reads (∼2-3%) presented a potentially artifactual methylation pattern wherein the 3’ end (but not the 5’ end) of these reads was demethylated (a pattern that could be explained by DNA nicks promoting 5’ to 3’ strand-displacing DNA synthesis during the FFPE DNA repair/End prep step of library preparation), we conservatively opted to filter out methylation calls exhibiting this pattern. Accordingly, we began read-based analyses by “clipping” methylation information from 3’ terminal regions of reads that lack modified CpGs (newly synthesized DNA is unmodified). These clipped reads were then further trimmed to include only that portion of the read that overlapped our ROI. We retained those reads that (post clipping and trimming) cover at least 75% of the CpGs in ROI. Average methylation for each read was then computed, enabling us to compare proportions of reads that were methylated above/below a threshold between disease conditions using a binomial mixed-effects model, including sex and age as covariates. This analysis is distinguished from the CpG-position based analysis in that read-membership information is not thrown out, allowing us to resolve changes in methylation involving a few, very hypo/hyper methylated reads versus many, subtly hypo/hyper methylated reads. This distinction holds biological significance because the sequenced molecules are native DNA molecules (have not been amplified by PCR) and hence all reads overlapping a given region very likely derive from different cells and can inform an assessment of cell-cell heterogeneity. To visualize read-level methylation events, we used the wgbstools package[65].

### Genotyping and read phasing

Filtered BAMs were input into the PEPPER-Margin-DeepVariant pipeline[66], which uses recurrent neural networks to perform long read-based variant calling and a hidden Markov model (HMM) for read phasing. We first annotated the resulting high-confidence small variant VCFs with SnpEff[67] and then added dbSNP[68] annotations using Bcftools[60]. *APOE* genotype (ɛ2/ɛ2, ɛ2/ɛ3, ɛ2/ɛ4, ɛ3/ɛ3, ɛ3/ɛ4, or ɛ4/ɛ4) was manually called on the basis of rs429358 and rs7412 values for each sample (ɛ2: rs429358 T; rs7412 T, ɛ3: rs429358 T; rs7412 C, ɛ4: rs429358 C; rs7412 C). Five of our twelve samples had their *APOE* genotype reported by the NIH NeuroBioBank, and all results were concordant with our analysis.

### Non-reference germline insertion calling and RTE-patching sample genomes

We used our TE-Seq pipeline[32] to call non-reference RTE insertions and create sample-specific, non-reference TE-patched genomes. Full methods can be found at (biorxiv.org/content/10.1101/2024.10.11.617912v1). Briefly, we aligned reads to the T2T-HS1 genome using Minimap2[69] and used TLDR[31] to call non-reference insertions. Non-reference insertions were annotated as “known” or “novel” on the basis of having been identified in a database of 17 datasets of human variation; see Table S4 in Ref[31]). Putative insertions were then rigorously filtered. We required that the median MAPQ score of supporting reads be 60 (the maximum possible), that the insert have a TSD (a hallmark of retrotransposition), and that the insert have at least 10 supporting reads, three of which must fully span the insert. We then appended each insert consensus sequence, flanked by 4 kilobases (kb) of insert-site sequence to enable accurate mapping of long reads, to the T2T-HS1 genome as a standalone contig. We annotated these contigs for repetitive element content with RepeatMasker[70] and merged these annotations with the T2T-HS1 reference RepeatMasker annotations.

### Somatic L1HS/AluY insertion calling

Starting with the list of putative RTE insertions called by TLDR (see above), we selected those events that might be consistent with a somatic event, i.e., they are supported by so few reads as to be inconsistent with an undersampled heterozygous germline insertion. We loosely estimated the likelihood of miscalling a heterozygous non-reference germline insertion as somatic, given the numbers of supporting versus empty reads. We modeled the haplotype sampling process as a binomial distribution, where the number of successes corresponds to the number of reads supporting the insertion, the number of trials is equal to the total coverage at the insertion site, and the probability of success (i.e., sampling a read containing the insertion) is set to 0.5.

Insertions with a p-value greater than 0.001 were excluded from further analysis. This threshold ensures that the number of insertion-supporting reads is significantly lower than the number of empty reads; for example, at the p < 0.001 level, at least 13 empty reads are required to consider a single insertion-supporting read as consistent with a somatic event.

We then required that the putative somatic insert be an L1HS or AluY element and be fully spanned (read encompasses the entire insertion and flanking sequence on both sides) by at least one read and that the median MAPQ score of all supporting reads be 60 (highest possible). Next, we applied an extensive set of filters to guard against spurious calls resulting from diverse sources including alignment error, contamination, and non-TPRT mediated mutations (e.g., products of homologous recombination or tandem duplications) (Fig. S4).

We filtered out any insertions that were called as “known” by TLDR, i.e., ones that had been identified in human population genetic variation studies (see Table S4 in [31] for more information on the 17 datasets comprising this database), as well as all insertions that were detected in more than one sample (be they somatic or germline). The rationale here is that a true somatic event is almost certainly unique, never before reported as a germline insertion or shared among samples. If an insertion is shared or a known germline mutation it is more likely this is either a sampling error (allelic bias), an alignment error, or contamination.

We then applied a set of insert-sequence-based filters. We required that the insert have a target site duplication (a hallmark of TPRT) of 21 bp or less. The insert sequence identity to the RTE consensus sequence had to be of 90% or more. Finally, since TPRT initiates from the 3’ end of the RTE, the 3’ end of the insert had to surpass the 250^th^ bp in the AluY consensus sequence or the 5,800^th^ bp in the L1HS consensus sequence (Fig. S4B).

To rein in spurious insertion calls derived from improperly aligned reads, we applied a set of spanning read and insert-site filters. All spanning reads had to be primary alignments and must not have any associated supplementary alignments. Furthermore, these reads must not have any indels or soft/hard clips greater than 50 bp in length. We required that the average MAPQ for all alignments at the insert site be greater than 55 and that the insert site itself be uniquely mappable by a 50 bp read. We flagged any insert sites that occurred in telomeres, centromeres, or segmental duplications for enhanced manual scrutiny.

We generated several plots and sequence files to aid in manual inspection of each event. A dotplot showing homology between the read and insert site using a 10 bp sliding window with a 1 mismatch tolerance was created for the 1,000 bp flanking the insert (Fig. S4D). This plot revealed if there was substantial homology between the inserted sequence and adjacent sequences. Such a situation could indicate that the event was a local, small duplication and was hence filtered out. To further guard against being misled by local duplications, we annotated the insert consensus sequence + 1,000 bp of insert site flanking sequence with RepeatMasker and verified that the inserted RTE was indeed detected and that no RTEs of the same subfamily were adjacent to the inserted sequence. We further ensured that the putative cut site had a motif consistent with ORF2’s cutting preference.

We next examined read phasing information derived from the PEPPER-Margin[66] pipeline in a genome browser. If insert bearing reads were assigned a haplotype (HP:1 or HP:2), we ensured that they did not exclusively comprise this haplotype (e.g., all reads with HP:1 had the insertion); this would suggest that the insertion might be an undersampled heterozygous germline variant. If insert-bearing reads were assigned to HP:0 (unphased), we verified that this was because there was insufficient information to distinguish between an HP:1 or HP:2 assignment, and not that the mutational profile was inconsistent with either haplotype.

Our final check filtered out false-positive somatic insertion calls derived from a chimeric read. Chimeric sequences are formed as an off-target reaction during library preparation when, instead of ligating sequencing adapters onto DNA, two DNA molecules are first ligated together. The vast majority of chimeric reads are readily identified by the Minimap2 aligner and have tell-tale supplementary alignments. In the unlikely event that a chimeric sequence is formed such that the shorter of the two concatenated DNA sequences contains an RTE followed by a stretch of sequence with high homology to the genomic sequence adjacent to the larger of the two concatenated DNA sequences, Minimap2 may fail to split the alignment into primary and supplementary alignments. It instead produces a single alignment with an insertion corresponding to the RTE and an imperfect downstream stretch of aligned sequence. To filter out such events, we split putative somatic insert bearing reads into (i) a 5’ sequence, which begins at the start of the read and terminates at the end of the putative insert, and (ii) a 3’ sequence that begins at the putative insert and extends until the end of the read. We then performed a nucleotide BLAST against the human genome and ensured that both sequence (i) and (ii) reported their best alignments at the putative insert site. Only after meeting all these filters did we call putative somatic L1HS/AluY insertions.

While the highly stringent filtering and manual inspection of events described above lends confidence to our analysis, we cannot exclude the possibility that some of our putative somatic events are false-positive calls attributable to either (i) misalignment of insert-containing reads; (ii) a loss of heterozygosity event at a germline RTE insertion paired with a clonal amplification of cells homozygous for the empty site; or (iii) contaminating DNA. These three scenarios could lead to a misleadingly low number of reads supporting what is in fact a germline insertion (which would then by virtue of the very low number of supporting reads be miscalled as a somatic insertion). The first of these concerns is attenuated by the fact that the misalignment of roughly half of the reads overlapping the insert site would be readily detectable as a loss of local coverage (similar to that observed in cases of heterozygous deletions), which we did not observe. The latter two concerns are mitigated by the fact that the vast majority of non-reference germline insertions are shared among individuals, and hence these events would be filtered out.

### Functional enrichment analysis of DMRs

We used the R Genomic Regions Enrichment of Annotations Tools (rGREAT) package[37] to find gene sets associated with DMRs. GREAT[71] models associations between genomic regions and gene sets in two ways, using either a binomial or a hypergeometric distribution. To increase biological interpretability, we examined only those DMRs that overlapped either gene promoters (−5 kb to +1 kb region flanking a RefSeq[72] gene transcription start site) or annotated enhancers (as identified by chromHMM). We used as background all gene promoters and enhancers. We queried Molecular Signatures Database (mSigDB)[73, 74] subcollections (e.g., GO:BP) for enrichment. For visualization, we ranked gene sets according to the mean of their binomial and hypergeometric FDR-adjusted p-value.

### RNA sequencing analysis

We performed stranded, paired-end 150 bp RNA-Seq for all samples on PFC, Brodmann area 9 tissue. Fresh frozen tissue was shipped to Azenta for library preparation and sequencing. RIN was measured using an Agilent RNA TapeStation. We used an rRNA depletion library preparation. Samples were comparable in terms of summary sequencing statistics, exhibiting high read quality, and mapping scores (Table S2). We analyzed these data using our TE-Seq pipeline. Full methods can be found at https://www.biorxiv.org/content/10.1101/2024.10.11.617912v1.

Briefly, sequencing reads were trimmed using fastp[75] and aligned to a sample-specific TE-patched genome (see Methods: *Non-reference Germline Insertion Calling and RTE-Patching Sample Genomes*) using STAR[76]. Gene expression was quantified using featureCounts[77] and a RefSeq transcript annotation (GCF_009914755.1). Repetitive element expression was quantified using the telescope[78] tool in “unique” mode, which only considers uniquely mapped reads. Genes were assessed for differential expression across disease condition using DESeq2[79], adjusting for RIN, age, and sex. We then used GSEA to query mSigDB gene sets for enrichment. Genome tracks were plotted using Plotgardener[80]. Differential expression analysis of aggregated RTE subfamily counts was performed using a negative binomial GLM. For this analysis, raw counts were summed across group members, and sample size factors estimated by DESeq2 were incorporated as an offset in the model. Disease condition, sex, age, and RIN were modeled as fixed effects.

Given that some RIN values were low (Table S2), we opted for an rRNA-depletion library preparation strategy. While this strategy avoids biasing sequencing against longer transcripts, it has as a downside that un-spliced pre-mRNA transcripts are retained for sequencing. As expected, a high fraction of our sequencing reads was intronic (Fig. S9C). This complicates RTE expression quantification because many RTEs reside in introns. When breaking down FL L1HS expression by genomic location (Fig. S9D), we found that intronic-sense elements had much higher expression values than did non-intronic-sense elements, suggesting passive expression through the sequencing of pre-mRNA introns. We therefore focused on the expression of non-intronic-sense elements. Furthermore, since RIN was associated with the fraction of reads mapping to exonic, intronic, and intergenic (though not repetitive element) regions (Fig. S9C), we used limma[81] to adjust DESeq2 normalized counts for the effect of RIN prior to downstream analysis of RTE counts. We restricted our analysis to uniquely mapping reads so as to prevent erroneously assigning intron-derived L1HS reads to non-intronic L1HS elements. However, this approach limited our ability to accurately quantify locus level L1HS expression, since many reads mapping to young L1 elements are non-uniquely mapped. Consequently, we focused on subfamily level expression estimates, which are more robust when only considering uniquely mapped reads.

## Declarations

### Ethics approval and consent to participate

For human brain tissue, written informed consent for post-mortem brain donation was obtained from the families of donors through the Mount Sinai Brain Bank (NIH NeuroBioBank). Prior to their transfer to the Icahn School of Medicine at Mount Sinai, all samples were de-identified, thereby exempting them from the oversight of the Institutional Review Board (IRB).

### Consent for publication

Not applicable

### Availability of data and materials

A Zenodo archive of all code used can be found at: 10.5281/zenodo.17805875. The most up-to-date version of the TE-Seq pipeline (which was used for sample-specific, non-reference TE-patched genome generation and RNA-seq analysis) can be found at https://github.com/maxfieldk/TE-Seq, where instructions for pipeline installation, configuration, and deployment are documented. All sequencing data have been deposited in the Sequencing Read Archive (SRA); BioProject accession number PRJNA1376503 (PFC RNA-Seq), PRJNA1376494 (PFC Nanopore DNA-Seq), PRJNA1375768 (LF1 Nanopore DNA-Seq) and PRJNA1376496 (N2102Ep Nanopore DNA-Seq). Any other data or information relevant to this study are available from the corresponding author upon reasonable request.

## Authors’ contributions

Conceptualization: ACP, AS, JL, JMS, MST, VG

Data curation: AC, MMK

Formal analysis: MMK

Funding acquisition: ACP, JL, JMS

Investigation: MMK, JMS

Methodology: AC, ACP, FHG, JDB, JMS, MMK

Project administration: ACP, JMS

Resources: ACP, JMS

Software: MMK

Supervision: ACP, AS, FHG, GDB, JL, JMS, MST, VG

Writing – original draft: MMK, JMS

Writing—review & editing: All authors

## Acknowledgments

The MMK and JMS would like to thank Jess Anderson for lab management and technical assistance, and members of the Sedivy lab and the Center on the Biology of Aging for feedback and support. We are grateful to the Brown Center for Computation and Visualization (CCV) team for their management of the OSCAR high performance computing cluster which was used throughout this work.

---

**Fig. S1.**
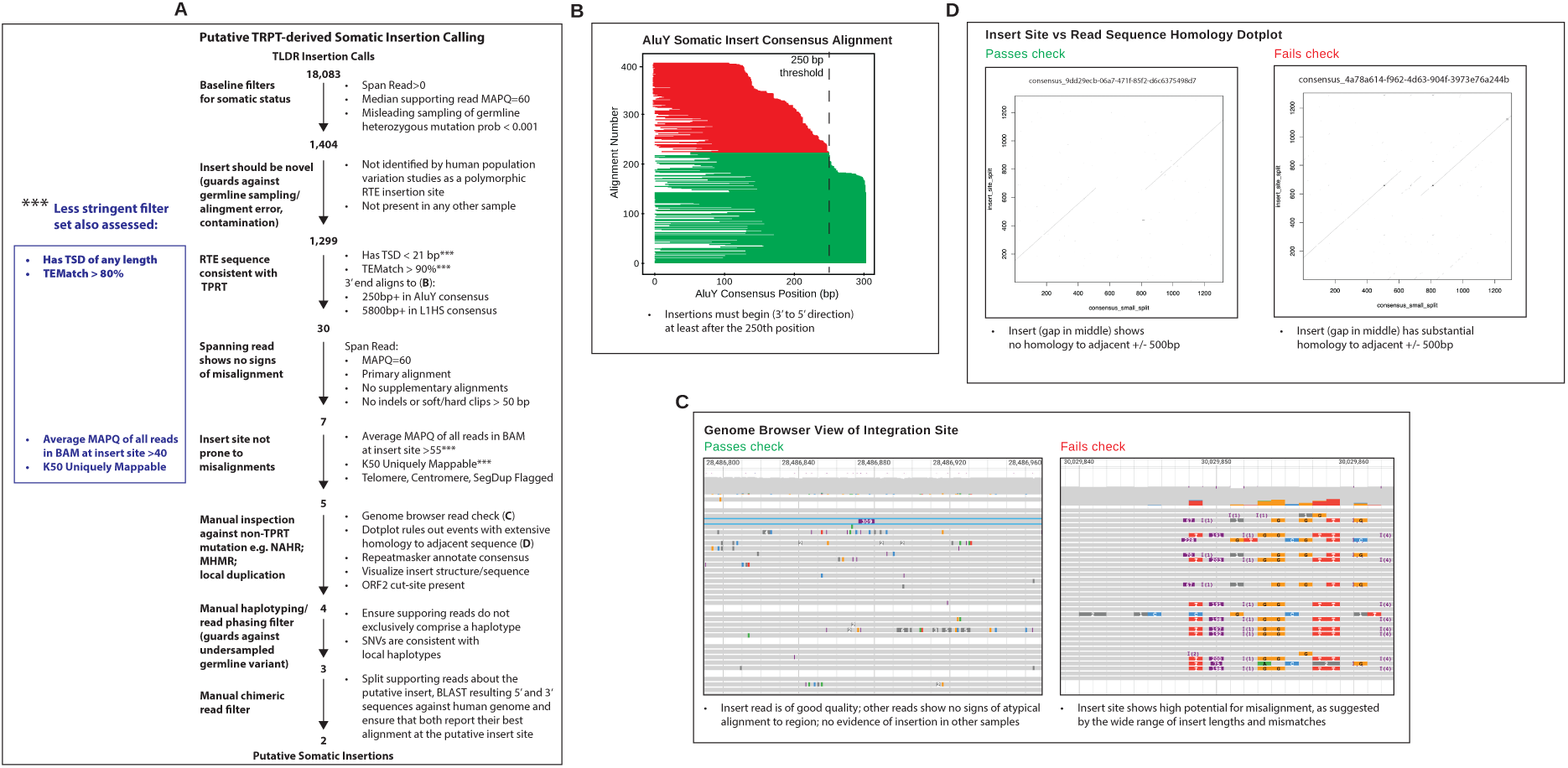
(**A**) TPRT-derived somatic insertion filtering strategy. (**B**) Somatic insertion 3’end filtering step visualized: AluY insertions whose 3’end occurs before the 250^th^ bp in the AluY consensus sequence (5800^th^ bp for L1HS) are filtered out. Red insertions (∼180) are filtered out, and green insertions (∼220) are retained. (**C**) Genome browser view of all alignments at two putative somatic insert sites. Large insertions are represented as purple boxes with the inserted bp count in white. The insert site on the left passes manual inspection, whereas the insert site on the right does not (there is potential read misalignment as suggested by the wide range of insert lengths shown). (**D**) Dot plot showing homology between the insert site (y-axis) and the read (x-axis) sequences. The insert on the left shows the expected linear tracking of reference sequence, followed by a ∼300 bp insertion of novel (non-homologous to insert region) sequence, which is, in turn, followed by the resumption of reference tracking (shifted down ∼20 bp owing to the TSD). The insert on the left shows extensive homology to the surrounding sequence and is not consistent with TPRT.

**Fig. S2.**
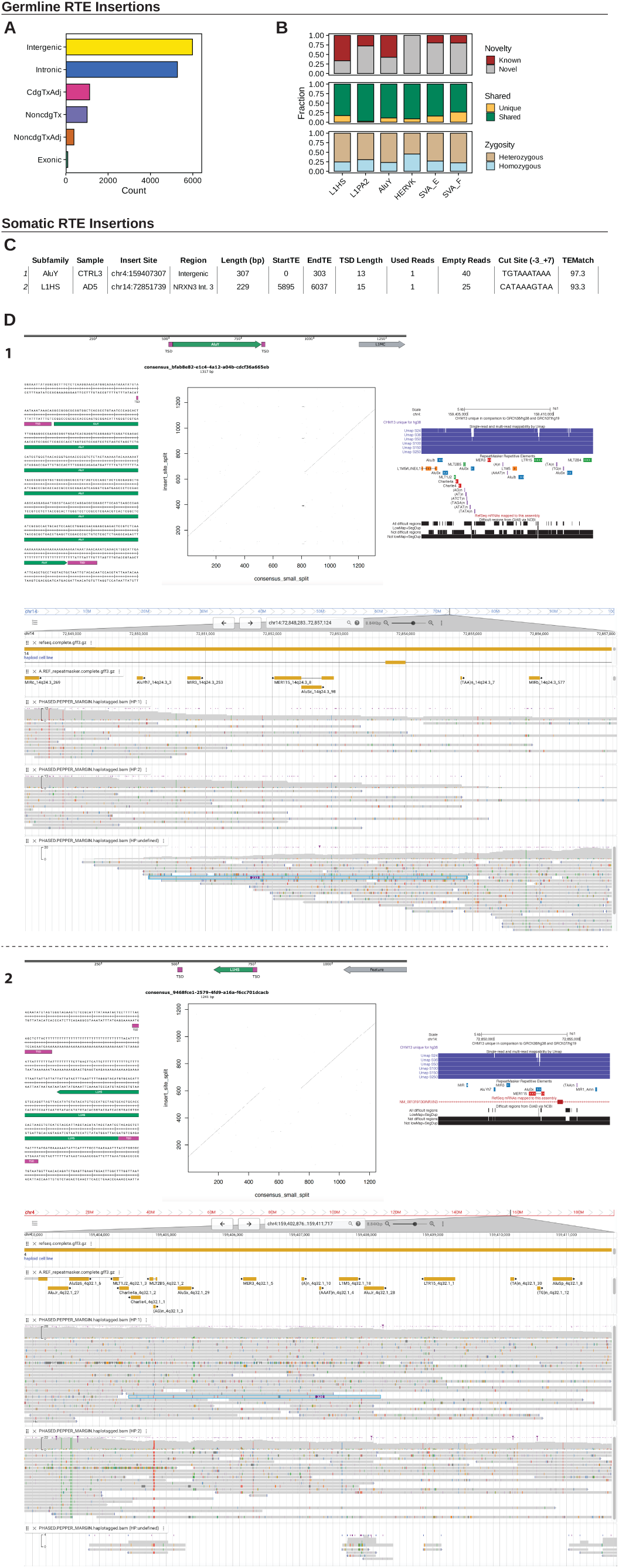
Evidence supporting somatic insertion calls in the aged brain. (**A**) Germline insertion counts grouped by their positional relationships to cellular genes (5’ and 3’ UTRs are included in the exonic category; Tx, transcript; Adj, adjacent: +/- 10kb from RefSeq curated transcript). (**B**) Bar graph depicting the fraction of germline insertions across all samples that are known/novel (top), shared/unique (middle), homozygous/heterozygous (bottom). Insertions denoted as ‘known’ have been previously identified in human populations, and insertions that are denoted as ‘shared’ were identified in more than one sample. (**C**) Table detailing putative somatic insertions. (**D**) Evidence supporting putative somatic insertion calls. For each insert listed in (**C**) we show (top) a SnapGene diagram of the insert + 1000bp of flanking sequence, annotated by RepeatMasker for repetitive element content; (left) a SnapGene representation of the inserted sequence; (middle) a dot plot showing homology between the insert site (y-axis) and the read (x-axis) sequences; (bottom) a genome browser view of phased (HP:1, HP:2, or HP:undefined) sequencing read alignments at the putative somatic insert site. Large insertions are represented as purple boxes with the inserted bp count in white; (right) a UCSC genome browser view centered on the insert site.

**Fig. S3.**
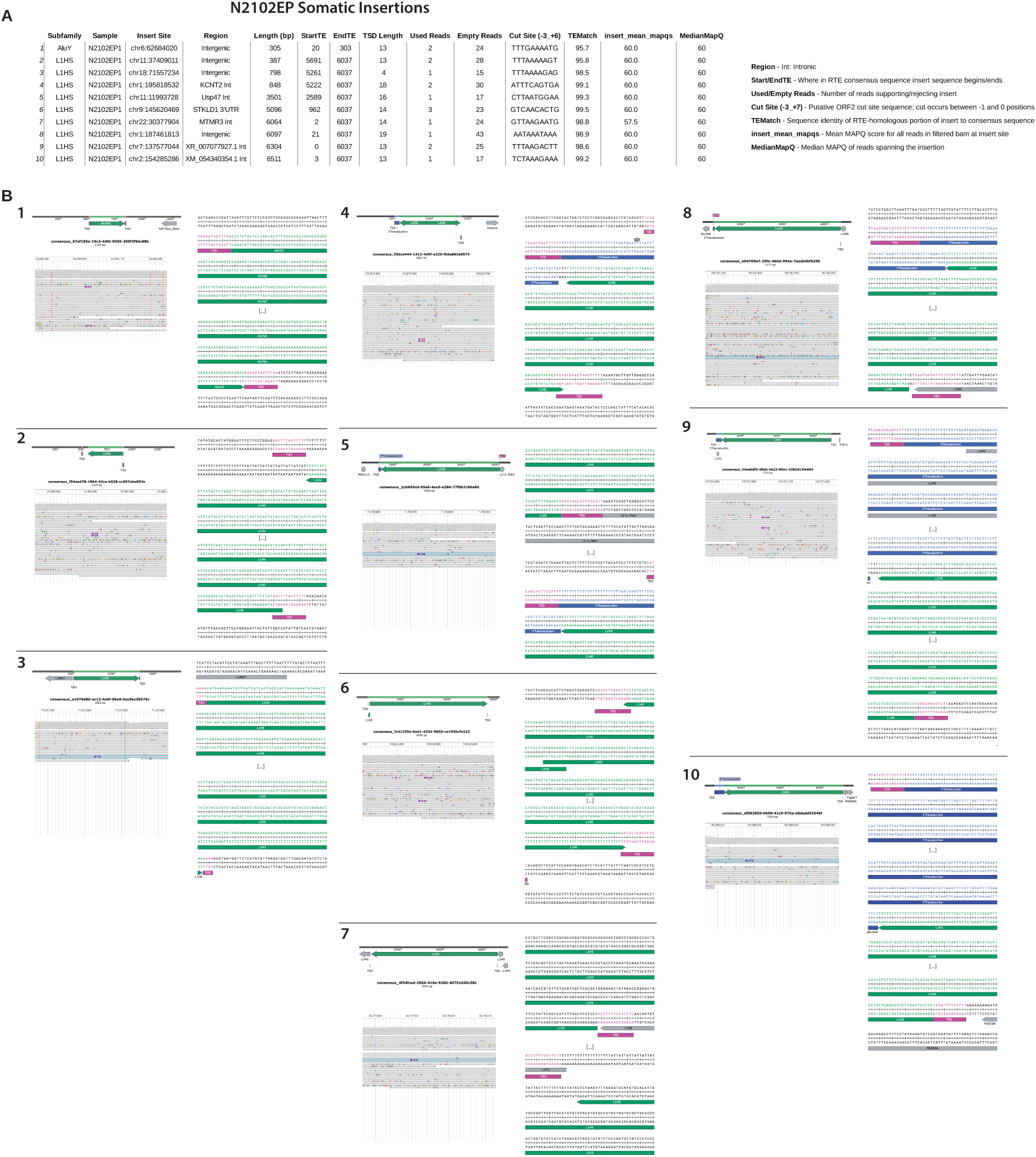
Evidence supporting somatic insertion calls in the N2102Ep cell line. (**A**) Table detailing putative somatic insertions in the N2102Ep cell line. (**B**) Evidence supporting putative somatic insertion calls. For each insert listed in (**A**) we show (top) a SnapGene diagram of the insert + 1000bp of flanking sequence, annotated by RepeatMasker for repetitive element content; (right) a SnapGene representation of the inserted sequence; (left) a genome browser view of all alignments at the putative somatic insert site. Large insertions are represented as purple boxes with the inserted bp count in white.

**Fig. S4.**
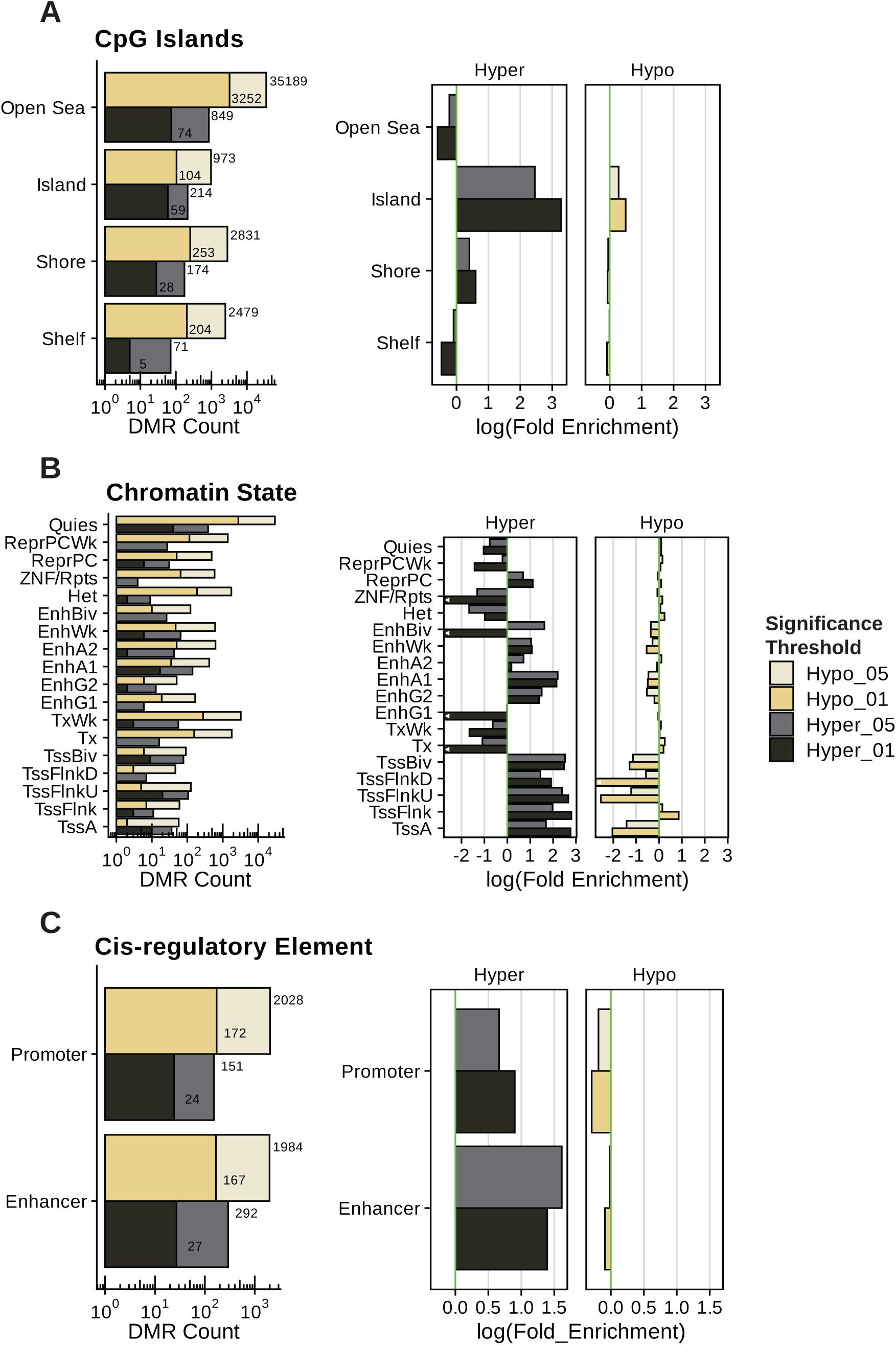
DMR overlaps and enrichments. We modeled the expected fraction of DMRs occurring in a set of regions as a binomial with probability equal to the genome fraction occupied by these regions: (**A**) CpG islands, (**B**) chromHMM states, (**C**) RefSeq gene promoters (−5 kb to +1 kb relative to TSS) and chromHMM-defined enhancers. For all panels, DMRs are colored by significance threshold and direction of change relative to CTRL. (**A-C**, left) Bar plots showing the number of DMRs that overlap a set of regions. (**A-C**, right) Bar plot of the natural logarithm of the fold enrichment of DMRs in these regions. Bars with a white arrow pointing left and touching the y-axis go to negative infinity. Quies: Quiescent/low, ReprPCWk: Weak repressed Polycomb, ReprPC: Repressed Polycomb, ZNF/Rpts: ZNF genes & repeats, Het: Heterochromatin, EnhBiv: Bivalent enhancer, EnhWk: Weak enhancer, EnhA2: Active enhancer 2, EnhA1: Active enhancer 1, EnhG2: Genic enhancer 2, EnhG1: Genic enhancer 1, TxWk: Weak transcription, Tx: Strong transcription, TssBiv: Bivalent/poised TSS, TssFlnkD: Flanking TSS downstream, TssFlnkU: Flanking TSS upstream, TssFlnk: Flanking TSS, TssA: Active TSS.

**Fig. S5.**
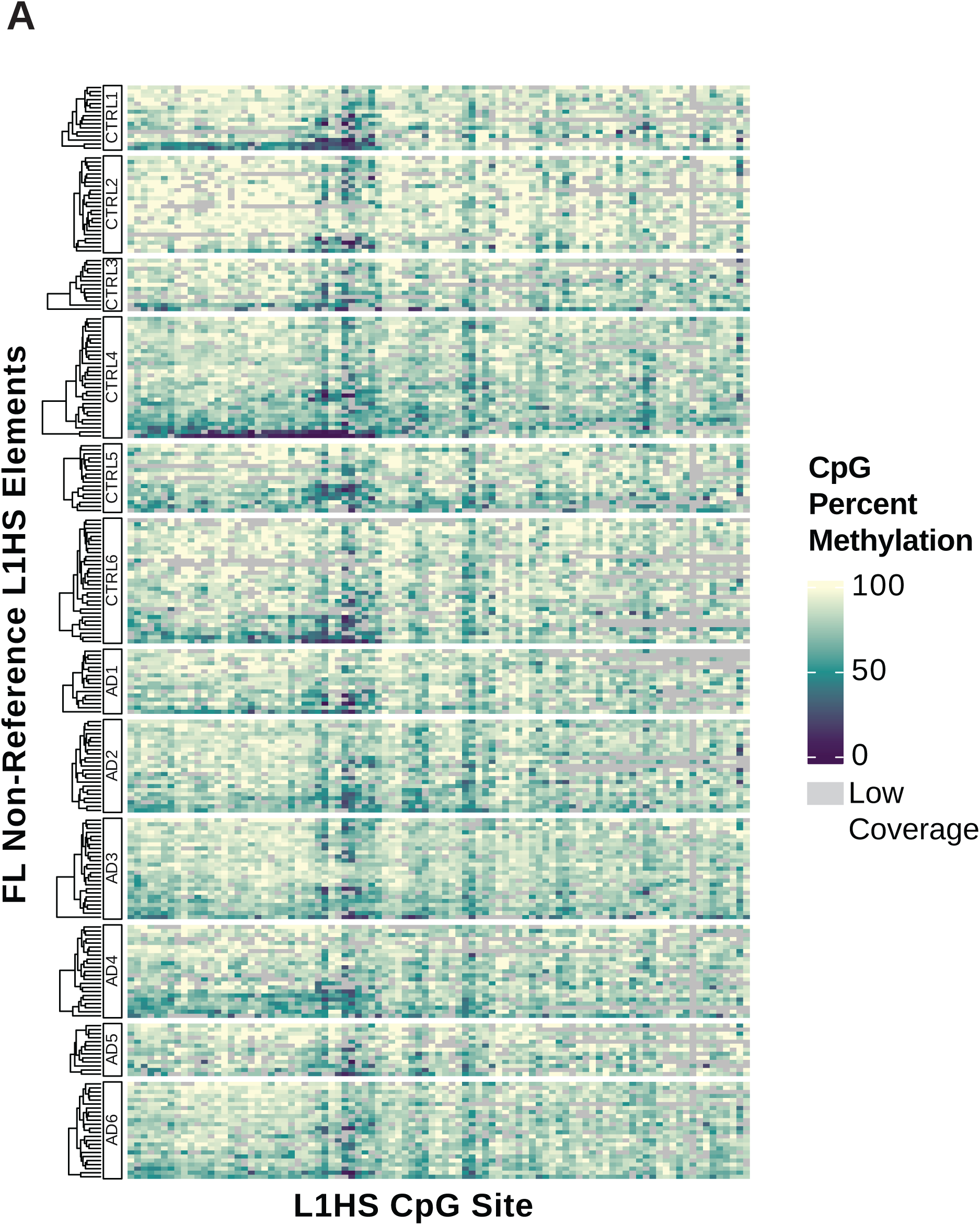
(**A**) Heatmaps of the mean CpG methylation for all non-reference FL L1HS elements. Heatmap rows represent individual L1HS elements and are hierarchically clustered. Heatmap columns represent individual CpG sites and are ordered by sequence position.

**Fig. S6.**
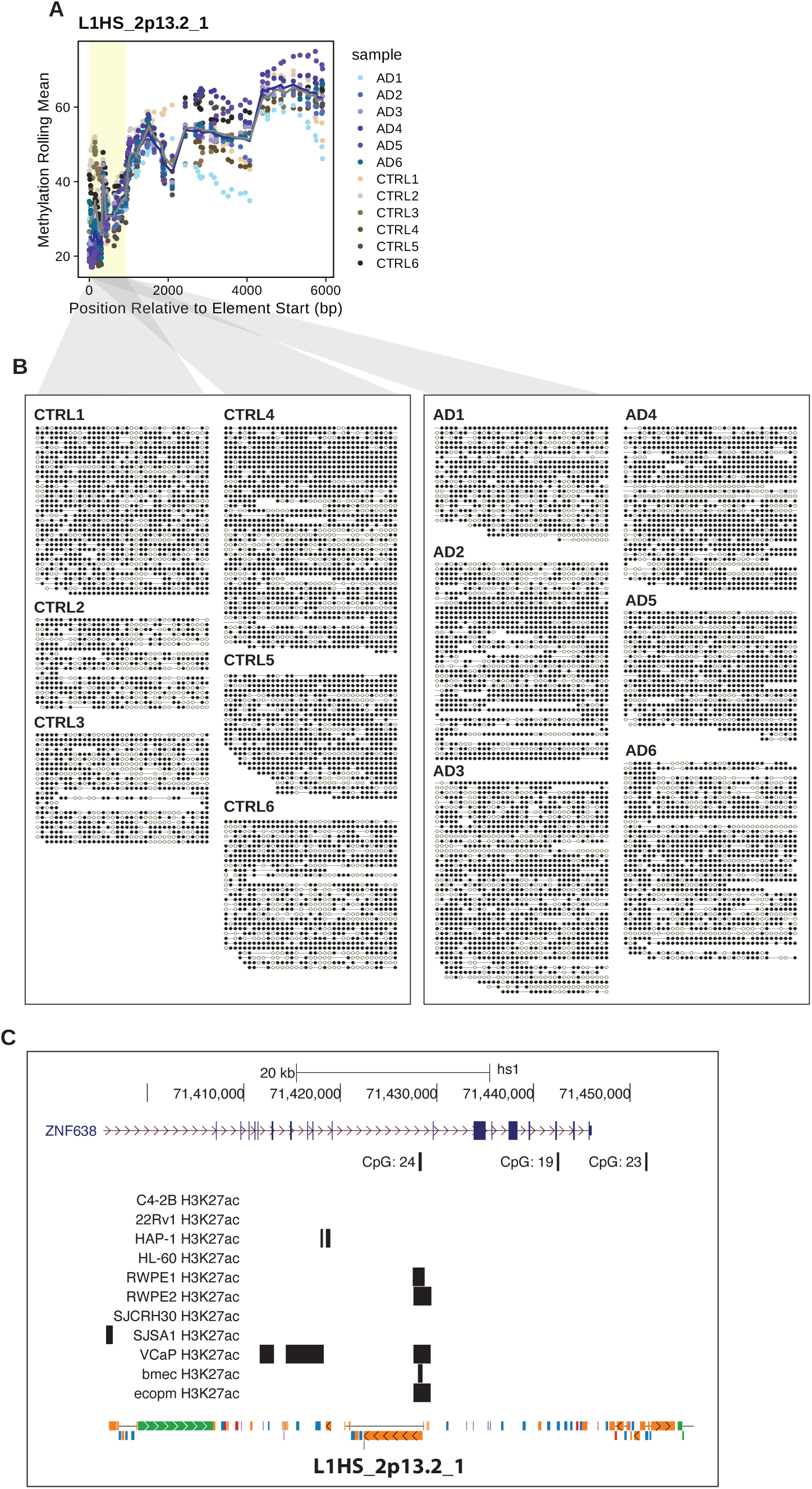
(**A**) Scatter plot showing the average methylation level by CpG for all CpGs in the L1HS_2p13.2_1 element. The 5’ UTR region is shaded in yellow. (**B**) Bubble plot showing all reads overlapping the L1HS_2p13.2_1 element’s 5’ UTR (region shaded in yellow). Reads are trimmed to not extend beyond this region. Each line segment represents a single sequencing read and each bubble represents a CpG site. The CpG is methylated if yellow, and demethylated if black. (**C**) UCSC genome browser view centered at the L1HS_2p13.2_1 element. From top to bottom we plot, RefSeq, CpG Island, ENCODE H3K27ac, and RepeatMasker tracks. Some labels are omitted for visual clarity. Several cell types show evidence of H3K27ac deposition at the L1HS_2p13.2_1 promoter.

**Fig. S7.**
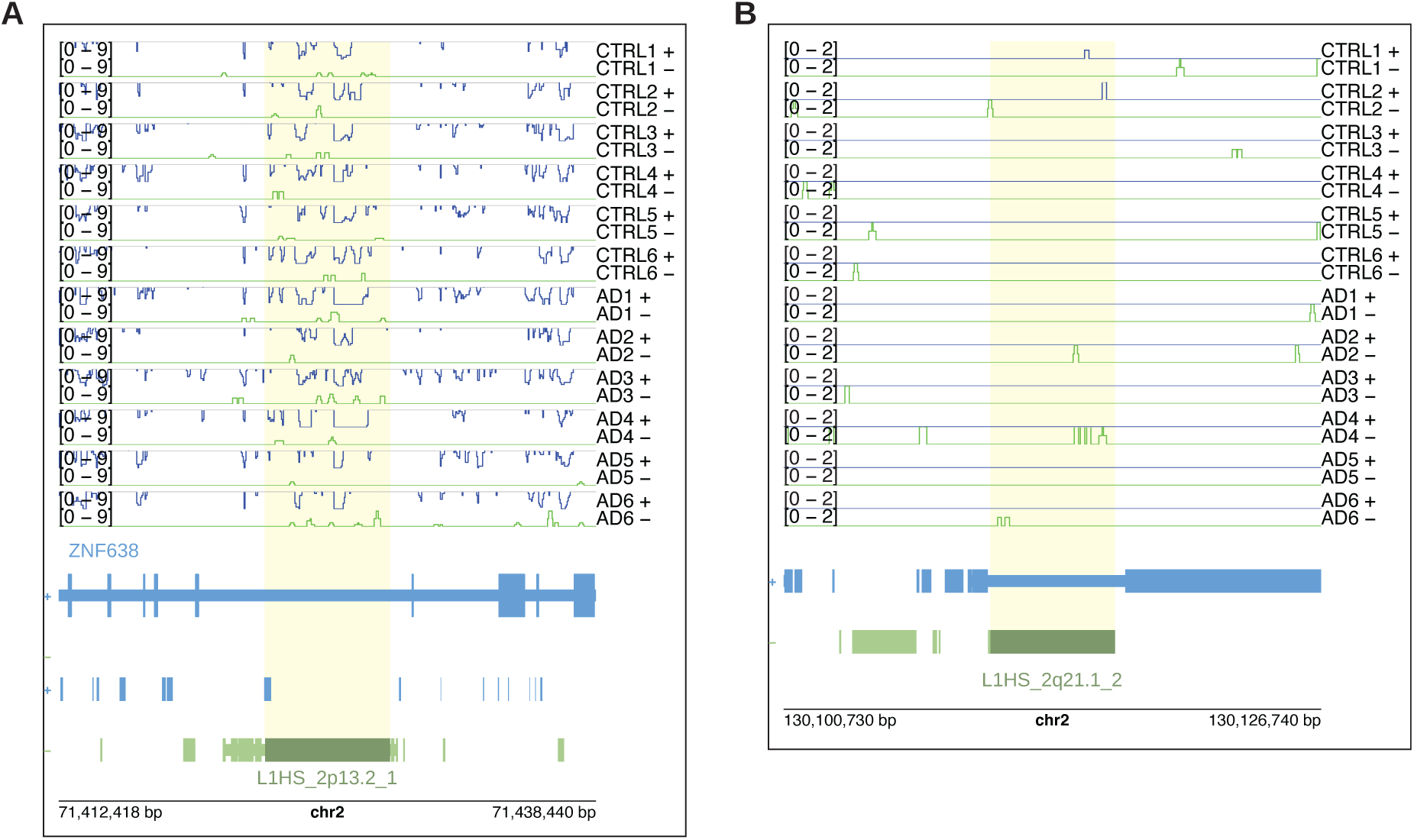
Locus Level L1HS RNA Expression. Genome tracks showing L1HS_2p13.2_1 (**A**) and L1HS_2q21.1_2 (**B**) expression. Primary alignments (secondary alignments are not shown) were converted to BigWig format and visualized in R using plotgardener. BigWig tracks of Watson (+; blue) and Crick (−; green) strand transcription are plotted for each sample, followed by RefSeq and RepeatMasker annotations. The region is centered on the L1HS element (boxed in black), with 10 kb of flanking up/downstream sequence. The BigWig signal track’s Y-axis upper bound is set at 110% the maximum count value observed across all samples over the L1HS locus.

**Fig. S8.**
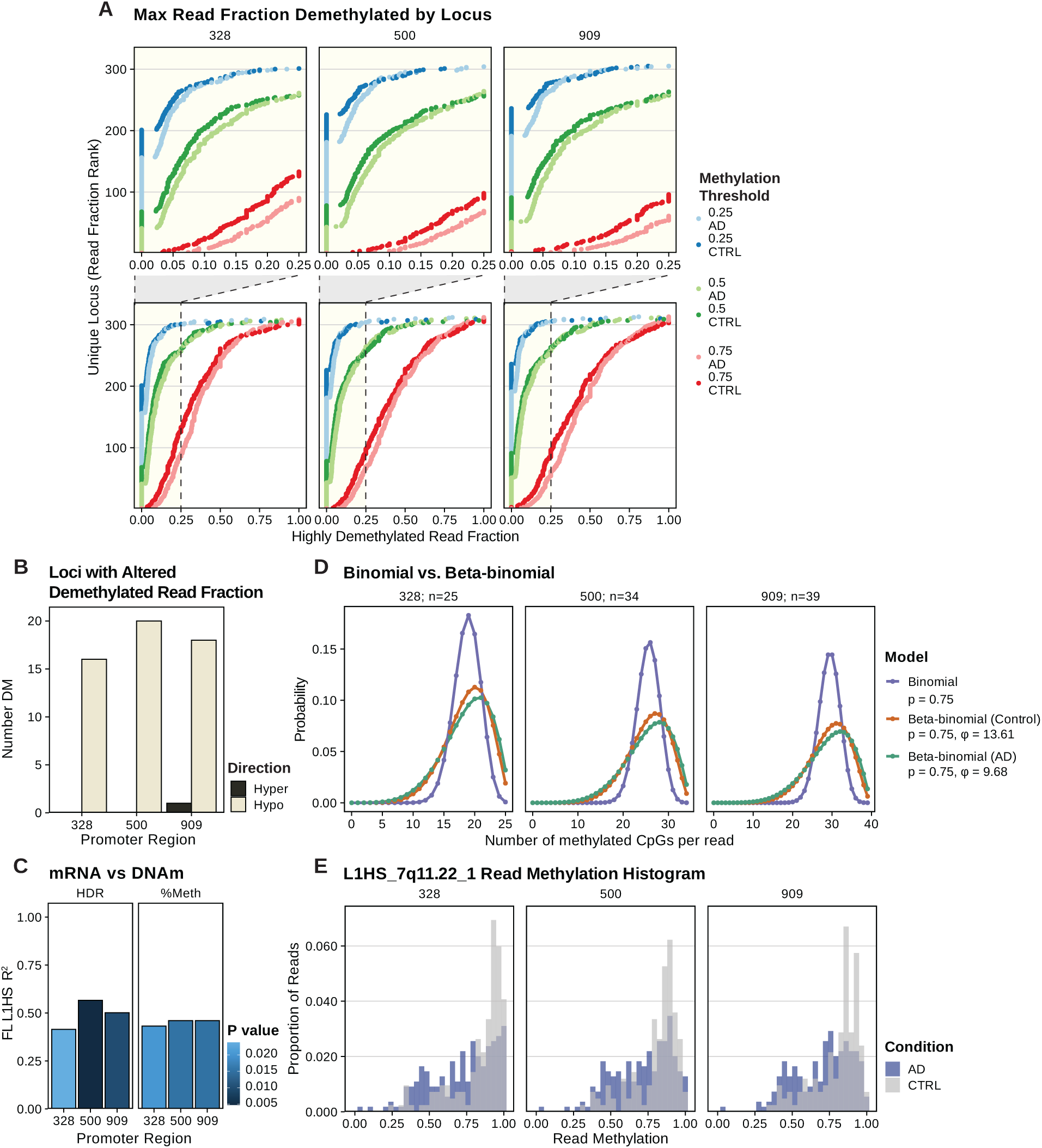
Full-length L1HS Read Level Methylation. (**A**) Dot plot showing, for each FL L1HS locus and condition, the sample with the greatest fraction of HDRs (colored by 0.25, 0.5, and 0.75 methylation thresholds) over the first 328 bp, 500 bp, 909 bp. (**B**) Bar plot showing the number of FL L1HS loci that have a significantly increased/decreased number of HDRs. Significance was assessed for each element using a binomial mixed-effects model as in **B**, with *p* ≤ 0.05. (**C**) We performed stranded, paired-end RNA-seq on rRNA-depleted RNA extracted from the same brain region used for Nanopore DNA-seq in each of our samples. We quantified RTE expression using uniquely mapped reads and the Telescope tool. To address confounding by intron retention, only RTE elements that do not reside in the sense orientation in a known transcript (RefSeq + Gencode) were used. We plot R^2^ values for the linear models fitting FL L1HS sense promoter expression as a function of DNA methylation for various 5’ UTR regions. (**D**) Probability mass functions (PMFs) illustrating the effect of overdispersion on read-level methylation. Each curve shows the expected distribution of methylated CpGs per read under a beta–binomial model with mean methylation p = 0.75 and different precision (φ) values: φ → ∞ (equivalent to a binomial), φ = 13.61 (estimated for control samples), and φ = 9.68 (estimated for AD samples). Lower φ corresponds to greater overdispersion and produces a wider, heavier-tailed distribution. (**E**) Histogram showing the distribution of read methylation over regions in the L1HS_7q11.22_1 5’ UTR.

**Fig. S9.**
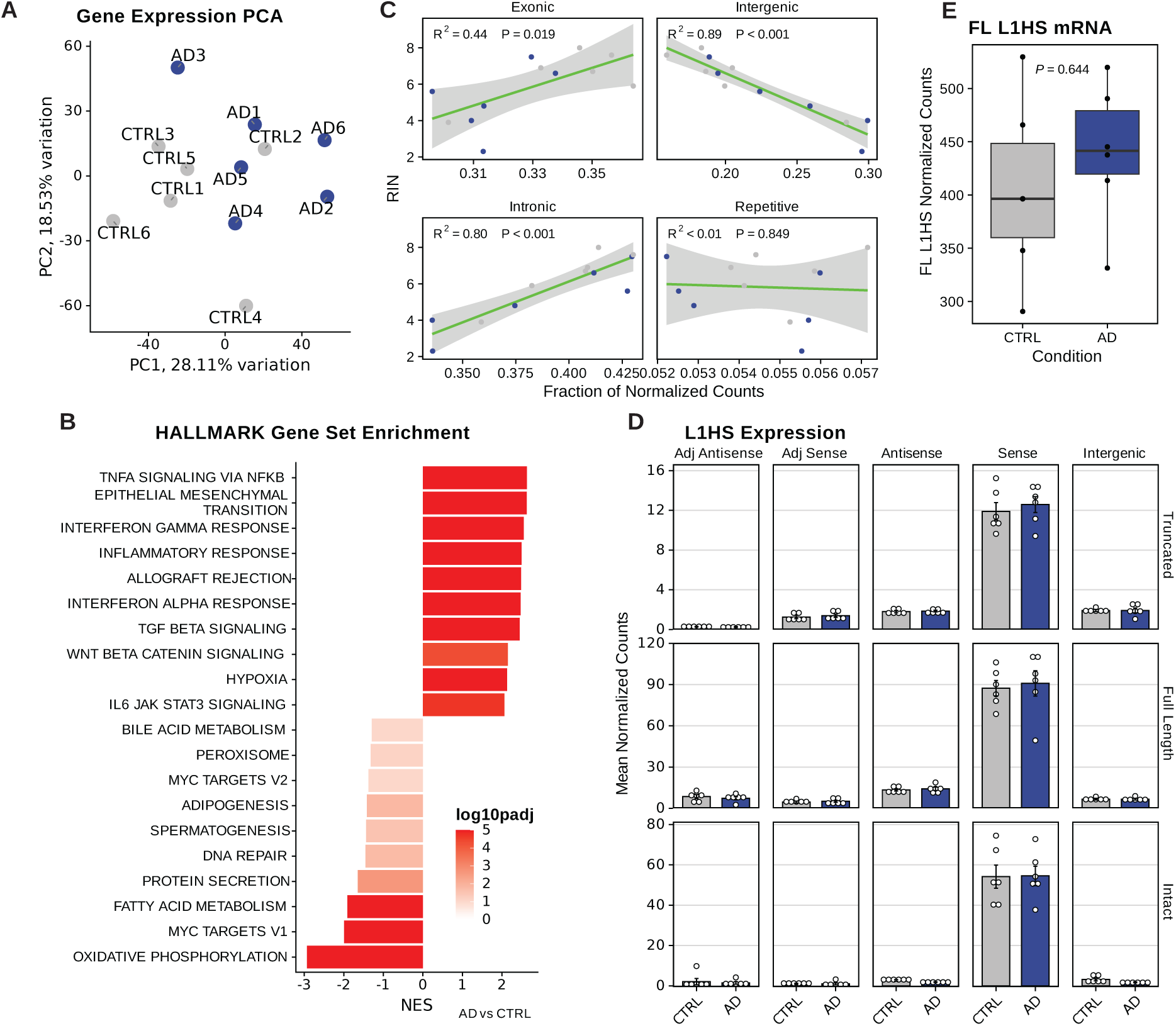
RNA-Seq. We performed stranded, paired-end RNA sequencing on rRNA-depleted RNA extracted from the same brain region used for Nanopore DNA-Seq in each of our samples. We analyzed these data using our TE-Seq pipeline (see Methods), which enables expression quantification at reference and non-reference RTE elements. (**A**) PCA-biplot showing the first two principal components of gene expression. Samples are colored by disease condition. (**B**) Gene set enrichment analysis (GSEA) of the mSigDB Hallmark collection. The top 10 most enriched/depleted gene sets are shown. (**C**) Scatter plots showing the correlation between RNA-integrity (RIN) and the fraction of reads mapping to various genomic regions. (**D**) Bar plots of total DESeq2 normalized read counts derived from various categories of L1HS elements. L1HS elements are grouped by their genomic location in relation to genes (columns) and by their length and intactness status (rows). (**E**) Sum of DESeq2 normalized counts for FL L1HS elements. Statistical significance was assessed using a negative binomial GLM with fixed effects for disease condition, age, sex, RIN; DESeq2-derived size factors were provided as an offset (*p* = 0.644).

**Fig. S10.**
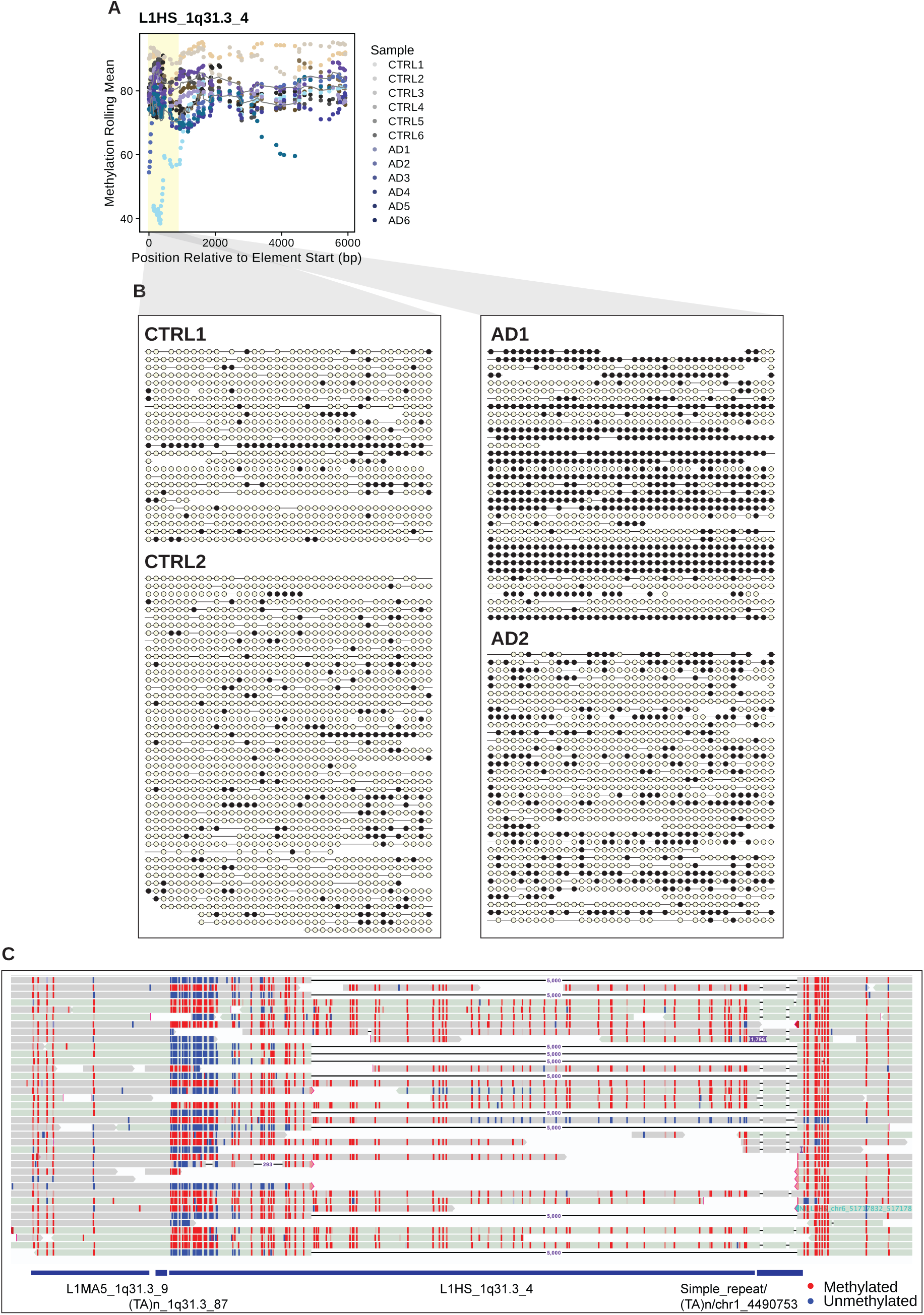
(**A**) Scatter plot showing the average methylation level by CpG for all CpGs in the L1HS_1q31.3_4 element. The 5’ UTR region is shaded in yellow. Condition averages are shown as lines. (**B**) Bubble plot showing all reads overlapping the L1HS_1q31.3_4 element’s 5’ UTR (region shaded in yellow). Reads are trimmed to not extend beyond this region. Each line segment represents a single sequencing read and each bubble represents a CpG site. The CpG is methylated if yellow, and demethylated if black. (**C**) IGV view of sample AD1 sequencing reads centered at the L1HS_1q31.3_4 element. Each line segment represents a single sequencing read. Reads are shaded by strand, and CpG sites are colored by methylation status (red: methylated, blue: unmethylated). Black line segments denote a 5 kb deletion which is found in roughly half of the sequencing reads. RepeatMasker annotations are found below sequencing reads.

**Table S1.**

|  | Sample | Sex | Age | Braak | APOE | Long read DNA-Seq |  |  |  | Short read RNA-Seq |  |
| --- | --- | --- | --- | --- | --- | --- | --- | --- | --- | --- | --- |
|  |  |  |  |  |  | Read Count | N50 | Basepair Count | Coverage | Read Count | RIN |
| 1 | CTRL1 | M | 79 | II | 2/3 | 1.95e+07 | 1.14e+04 | 9.76e+10 | 31.5 | 1.23e+08 | 8.0 |
| 2 | CTRL2 | M | 71 | III | 3/3 | 1.61e+07 | 1.14e+04 | 9.78e+10 | 31.5 | 9.92e+07 | 7.6 |
| 3 | CTRL3 | M | 81 | II | 3/3 | 2.15e+07 | 1.04e+04 | 1.00e+11 | 32.3 | 1.07e+08 | 6.7 |
| 4 | CTRL4 | F | 76 | II | 3/3 | 3.99e+07 | 6.48e+03 | 1.31e+11 | 42.4 | 1.01e+08 | 3.9 |
| 5 | CTRL5 | F | 88 | NA* | 3/3 | 1.85e+07 | 7.97e+03 | 7.31e+10 | 23.6 | 9.82e+07 | 6.9 |
| 6 | CTRL6 | F | 76 | IV | 3/4 | 1.21e+07 | 1.01e+04 | 8.43e+10 | 27.2 | 1.03e+08 | 5.9 |
| 7 | AD1 | F | 83 | VI | 3/4 | 2.57e+07 | 8.84e+03 | 1.09e+11 | 35.2 | 8.97e+07 | 5.6 |
| 8 | AD2 | M | 77 | V | 4/4 | 4.36e+07 | 5.61e+03 | 1.32e+11 | 42.6 | 1.18e+08 | 4.8 |
| 9 | AD3 | F | 70 | VI | 3/3 | 3.92e+07 | 5.95e+03 | 1.40e+11 | 45.2 | 9.77e+07 | 2.3 |
| 10 | AD4 | F | 89 | VI | 3/4 | 2.78e+07 | 9.41e+03 | 1.10e+11 | 35.3 | 1.08e+08 | 6.6 |
| 11 | AD5 | M | 74 | VI | 3/3 | 1.61e+07 | 8.80e+03 | 6.53e+10 | 21.1 | 9.58e+07 | 7.5 |
| 12 | AD6 | M | 74 | VI | 3/3 | 3.90e+07 | 6.27e+03 | 1.28e+11 | 41.2 | 1.15e+08 | 4.0 |

\*See sample selection in **methods**

**Table S2.**
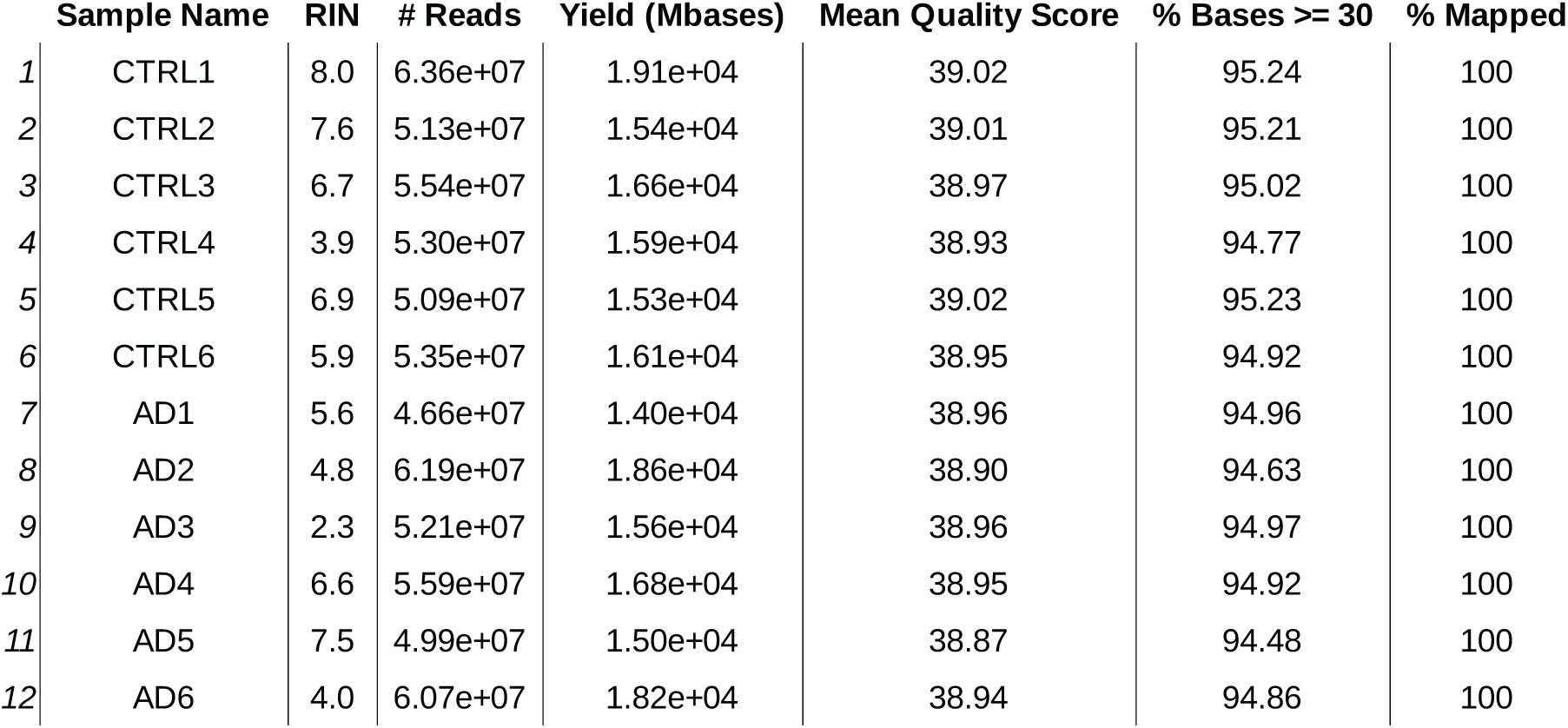

## References

1. DeTure MA, Dickson DW: The neuropathological diagnosis of Alzheimer’s disease. Mol Neurodegener 2019, 14:32.

2. Heneka MT, Carson MJ, El Khoury J, Landreth GE, Brosseron F, Feinstein DL, Jacobs AH, Wyss-Coray T, Vitorica J, Ransohoff RM, et al: Neuroinflammation in Alzheimer’s disease. Lancet Neurology 2015, 14:388–405.

3. Ransohoff RM: How neuroinflammation contributes to neurodegeneration. Science 2016, 353:777–783.

4. Hou Y, Dan X, Babbar M, Wei Y, Hasselbalch SG, Croteau DL, Bohr VA: Ageing as a risk factor for neurodegenerative disease. Nat Rev Neurol 2019, 15:565–581.

5. Pal S, Tyler JK: Epigenetics and aging. Sci Adv 2016, 2:e1600584.

6. Lu T, Pan Y, Kao SY, Li C, Kohane I, Chan J, Yankner BA: Gene regulation and DNA damage in the ageing human brain. Nature 2004, 429:883–891.

7. Bollati V, Schwartz J, Wright R, Litonjua A, Tarantini L, Suh H, Sparrow D, Vokonas P, Baccarelli A: Decline in genomic DNA methylation through aging in a cohort of elderly subjects. Mech Ageing Dev 2009, 130:234–239.

8. Heyn H, Li N, Ferreira HJ, Moran S, Pisano DG, Gomez A, Diez J, Sanchez-Mut JV, Setien F, Carmona FJ, et al: Distinct DNA methylomes of newborns and centenarians. Proc Natl Acad Sci U S A 2012, 109:10522–10527.

9. Lee JH, Kim EW, Croteau DL, Bohr VA: Heterochromatin: an epigenetic point of view in aging. Experimental and Molecular Medicine 2020, 52:1466–1474.

10. Gorbunova V, Seluanov A, Mita P, McKerrow W, Fenyö D, Boeke JD, Linker SB, Gage FH, Kreiling JA, Petrashen AP, et al: The role of retrotransposable elements in ageing and age-associated diseases. Nature 2021, 596:43–53.

11. Sulc P, Di Gioacchino A, Solovyov A, Sun S, Martis S, Marhon SA, Lindholm HT, Chen R, Hosseini A, Jiang H, et al: Repeats mimic pathogen-associated patterns across a vast evolutionary landscape. Cell Genom 2025:101011.

12. Bourque G, Burns KH, Gehring M, Gorbunova V, Seluanov A, Hammell M, Imbeault M, Izsvák Z, Levin HL, Macfarlan TS, et al: Ten things you should know about transposable elements. Genome Biology 2018, 19:199.

13. Gasior SL, Wakeman TP, Xu B, Deininger PL: The Human LINE-1 Retrotransposon Creates DNA Double-strand Breaks. Journal of Molecular Biology 2006, 357:1383–1393.

14. Deininger PL, Moran JV, Batzer MA, Kazazian HH: Mobile elements and mammalian genome evolution. Current Opinion in Genetics & Development 2003, 13:651–658.

15. Han K, Lee J, Meyer TJ, Remedios P, Goodwin L, Batzer MA: L1 recombination-associated deletions generate human genomic variation. Proceedings of the National Academy of Sciences of the United States of America 2008, 105:19366–19371.

16. De Cecco M, Ito T, Petrashen AP, Elias AE, Skvir NJ, Criscione SW, Caligiana A, Brocculi G, Adney EM, Boeke JD, et al: L1 drives IFN in senescent cells and promotes age-associated inflammation. Nature 2019, 566:73-+.

17. Alexandrova EA, Olovnikov IA, Malakhova GV, Zabolotneva AA, Suntsova MV, Dmitriev SE, Buzdin AA: Sense transcripts originated from an internal part of the human retrotransposon LINE-1 5′ UTR. Gene 2012, 511:46–53.

18. Sanchez-Luque FJ, Kempen M-JHC, Gerdes P, Vargas-Landin DB, Richardson SR, Troskie R-L, Jesuadian JS, Cheetham SW, Carreira PE, Salvador-Palomeque C, et al: LINE-1 Evasion of Epigenetic Repression in Humans. Molecular Cell 2019, 75:590–604.e512.

19. Deniz Ö, Frost JM, Branco MR: Regulation of transposable elements by DNA modifications. Nature Reviews Genetics 2019, 20:417–431.

20. Protasova MS, Andreeva TV, Rogaev EI: Factors Regulating the Activity of LINE1 Retrotransposons. Genes 2021, 12:1562.

21. Van Meter M, Kashyap M, Rezazadeh S, Geneva AJ, Morello TD, Seluanov A, Gorbunova V: SIRT6 represses LINE1 retrotransposons by ribosylating KAP1 but this repression fails with stress and age. Nature Communications 2014, 5:5011.

22. Simon M, Meter MV, Ablaeva J, Ke Z, Gonzalez RS, Taguchi T, Cecco MD, Leonova KI, Kogan V, Helfand SL, et al: LINE1 Derepression in Aged Wild-Type and SIRT6-Deficient Mice Drives Inflammation. Cell Metabolism 2019, 29:871–885.e875.

23. Mastroeni D, Grover A, Delvaux E, Whiteside C, Coleman PD, Rogers J: Epigenetic changes in Alzheimer’s disease: decrements in DNA methylation. Neurobiology of Aging 2010, 31:2025–2037.

24. Chouliaras L, Mastroeni D, Delvaux E, Grover A, Kenis G, Hof PR, Steinbusch HWM, Coleman PD, Rutten BPF, van den Hove DLA: Consistent decrease in global DNA methylation and hydroxymethylation in the hippocampus of Alzheimer’s disease patients. Neurobiology of Aging 2013, 34:2091–2099.

25. De Plano LM, Saitta A, Oddo S, Caccamo A: Epigenetic Changes in Alzheimer’s Disease: DNA Methylation and Histone Modification. Cells 2024, 13:719.

26. Xiong X, James BT, Boix CA, Park YP, Galani K, Victor MB, Sun N, Hou L, Ho L-L, Mantero J, et al: Epigenomic dissection of Alzheimer’s disease pinpoints causal variants and reveals epigenome erosion. Cell 2023, 186:4422–4437.e4421.

27. Wang B-A, Jones JR, Zhou J, Tian W, Wu Y, Wang W, Berube P, Bartlett A, Castanon R, Nery JR, et al: Epigenome erosion in Alzheimer’s disease brain cells and induced neurons. bioRxiv; 2023.

28. Frost B, Hemberg M, Lewis J, Feany MB: Tau promotes neurodegeneration through global chromatin relaxation. Nature Neuroscience 2014, 17:357–366.

29. Sun W, Samimi H, Gamez M, Zare H, Frost B: Pathogenic tau-induced piRNA depletion promotes neuronal death through transposable element dysregulation in neurodegenerative tauopathies. Nature Neuroscience 2018, 21:1038–1048.

30. Ochoa E, Ramirez P, Gonzalez E, De Mange J, Ray WJ, Bieniek KF, Frost B: Pathogenic tau–induced transposable element–derived dsRNA drives neuroinflammation. Science Advances 2023, 9:eabq5423.

31. Ewing AD, Smits N, Sanchez-Luque FJ, Faivre J, Brennan PM, Richardson SR, Cheetham SW, Faulkner GJ: Nanopore Sequencing Enables Comprehensive Transposable Element Epigenomic Profiling. Molecular Cell 2020, 80:915–928.e915.

32. Kelsey MMG, Kalekar RL, Sedivy JM: TE-Seq: A Transposable Element Annotation and RNA-Seq Pipeline. bioRxiv 2025:2024.2010.2011.617912.

33. Kamal N, Jafari Khamirani H, Dara M, Dianatpour M: NRXN3 mutations cause developmental delay, movement disorder, and behavioral problems: CRISPR edited cells based WES results. Gene 2023, 867:147347.

34. Jurka J: Sequence patterns indicate an enzymatic involvement in integration of mammalian retroposons. Proceedings of the National Academy of Sciences of the United States of America 1997, 94:1872–1877.

35. Feng Q, Moran JV, Kazazian HH, Jr., Boeke JD: Human L1 retrotransposon encodes a conserved endonuclease required for retrotransposition. Cell 1996, 87:905–916.

36. Di Stefano LH, Saba LJ, Oghbaie M, Jiang H, McKerrow W, Benitez-Guijarro M, Taylor MS, LaCava J: Affinity-Based Interactome Analysis of Endogenous LINE-1 Macromolecules. Methods Mol Biol 2023, 2607:215–256.

37. Gu Z, Hübschmann D: rGREAT: an R/bioconductor package for functional enrichment on genomic regions. *Bioinformatics (Oxford*, England) 2023, 39:btac745.

38. Seczynska M, Lehner PJ: The sound of silence: mechanisms and implications of HUSH complex function. Trends in Genetics 2023, 39:251–267.

39. Wallis NJ, McClellan A, Morseburg A, Kentistou KA, Jamaluddin A, Dowsett GKC, Schofield E, Morros-Nuevo A, Saeed S, Lam BYH, et al: Canine genome-wide association study identifies DENND1B as an obesity gene in dogs and humans. Science 2025, 387:eads2145.

40. Pal D, Patel M, Boulet F, Sundarraj J, Grant OA, Branco MR, Basu S, Santos SDM, Zabet NR, Scaffidi P, Pradeepa MM: H4K16ac activates the transcription of transposable elements and contributes to their cis-regulatory function. Nat Struct Mol Biol 2023, 30:935–947.

41. Upton KR, Gerhardt DJ, Jesuadian JS, Richardson SR, Sánchez-Luque FJ, Bodea GO, Ewing AD, Salvador-Palomeque C, van der Knaap MS, Brennan PM, et al: Ubiquitous L1 Mosaicism in Hippocampal Neurons. Cell 2015, 161:228–239.

42. Evrony Gilad D, Cai X, Lee E, Hills LB, Elhosary PC, Lehmann Hillel S, Parker JJ, Atabay Kutay D, Gilmore Edward C, Poduri A, et al: Single-Neuron Sequencing Analysis of L1 Retrotransposition and Somatic Mutation in the Human Brain. Cell 2012, 151:483–496.

43. Erwin JA, Paquola ACM, Singer T, Gallina I, Novotny M, Quayle C, Bedrosian TA, Alves FIA, Butcher CR, Herdy JR, et al: L1-associated genomic regions are deleted in somatic cells of the healthy human brain. Nature Neuroscience 2016, 19:1583–1591.

44. Evrony GD, Lee E, Park PJ, Walsh CA: Resolving rates of mutation in the brain using single-neuron genomics. eLife 2016, 5:e12966.

45. Hata K, Sakaki Y: Identification of critical CpG sites for repression of L1 transcription by DNA methylation. Gene 1997, 189:227–234.

46. Jönsson ME, Ludvik Brattås P, Gustafsson C, Petri R, Yudovich D, Pircs K, Verschuere S, Madsen S, Hansson J, Larsson J, et al: Activation of neuronal genes via LINE-1 elements upon global DNA demethylation in human neural progenitors. Nature Communications 2019, 10.

47. Beam CR, Kaneshiro C, Jang JY, Reynolds CA, Pedersen NL, Gatz M: Differences Between Women and Men in Incidence Rates of Dementia and Alzheimer’s Disease. Journal of Alzheimer’s disease : JAD 2018, 64:1077–1083.

48. Lanciano S, Philippe C, Sarkar A, Pratella D, Domrane C, Doucet AJ, van Essen D, Saccani S, Ferry L, Defossez P-A, Cristofari G: Locus-level L1 DNA methylation profiling reveals the epigenetic and transcriptional interplay between L1s and their integration sites. Cell Genomics 2024, 4:100498.

49. Garza R, Atacho DAM, Adami A, Gerdes P, Vinod M, Hsieh P, Karlsson O, Horvath V, Johansson PA, Pandiloski N, et al: LINE-1 retrotransposons drive human neuronal transcriptome complexity and functional diversification. Sci Adv 2023, 9:eadh9543.

50. Nicodemus J, Liu CS, Ransom L, Tan V, Romanow W, Jimenez N, Chun J: Sequence diversity and encoded enzymatic differences of monocistronic L1 ORF2 mRNA variants in the aged normal and Alzheimer’s disease brain. J Neurosci 2025.

51. Dileep V, Boix CA, Mathys H, Marco A, Welch GM, Meharena HS, Loon A, Jeloka R, Peng Z, Bennett DA, et al: Neuronal DNA double-strand breaks lead to genome structural variations and 3D genome disruption in neurodegeneration. Cell 2023, 186:4404–4421 e4420.

52. Welch G, Tsai LH: Mechanisms of DNA damage-mediated neurotoxicity in neurodegenerative disease. EMBO Rep 2022, 23:e54217.

53. Welch GM, Boix CA, Schmauch E, Davila-Velderrain J, Victor MB, Dileep V, Bozzelli PL, Su Q, Cheng JD, Lee A, et al: Neurons burdened by DNA double-strand breaks incite microglia activation through antiviral-like signaling in neurodegeneration. Sci Adv 2022, 8:eabo4662.

54. Jensvold ZD, Flood JR, Christenson AE, Lewis PW: Interplay between Two Paralogous Human Silencing Hub (HuSH) Complexes in Regulating LINE-1 Element Silencing. Nat Commun 2024, 15:9492.

55. Seetharam D, Chandar J, Ramsoomair CK, Desgraves JF, Medina AA, Hudson AJ, Amidei A, Castro JR, Govindarajan V, Wang S, et al: Targeting ZNF638 activates antiviral immune responses and potentiates immune checkpoint inhibition in glioblastoma. bioRxiv 2024.

56. Zhu Y, Wang GZ, Cingoz O, Goff SP: NP220 mediates silencing of unintegrated retroviral DNA. Nature 2018, 564:278–282.

57. Wheelan SJ, Aizawa Y, Han JS, Boeke JD: Gene-breaking: a new paradigm for human retrotransposon-mediated gene evolution. Genome Res 2005, 15:1073–1078.

58. Nurk S, Koren S, Rhie A, Rautiainen M, Bzikadze A, Mikheenko A, Vollger MR, Altemose N, Uralsky L, Gershman A, et al: The complete sequence of a human genome. Science 2022, 376:44-+.

59. Leger A, Leonardi T: pycoQC, interactive quality control for Oxford Nanopore Sequencing. Journal of Open Source Software 2019, 4:1236.

60. Danecek P, Bonfield JK, Liddle J, Marshall J, Ohan V, Pollard MO, Whitwham A, Keane T, McCarthy SA, Davies RM, Li H: Twelve years of SAMtools and BCFtools. GigaScience 2021, 10:giab008.

61. Park Y, Wu H: Differential methylation analysis for BS-seq data under general experimental design. Bioinformatics 2016, 32:1446–1453.

62. Lawrence M, Huber W, Pagès H, Aboyoun P, Carlson M, Gentleman R, Morgan MT, Carey VJ: Software for Computing and Annotating Genomic Ranges. PLOS Computational Biology 2013, 9:e1003118.

63. Luo Y, Hitz BC, Gabdank I, Hilton JA, Kagda MS, Lam B, Myers Z, Sud P, Jou J, Lin K, et al: New developments on the Encyclopedia of DNA Elements (ENCODE) data portal. Nucleic Acids Research 2020, 48:D882–D889.

64. Hinrichs AS, Karolchik D, Baertsch R, Barber GP, Bejerano G, Clawson H, Diekhans M, Furey TS, Harte RA, Hsu F, et al: The UCSC Genome Browser Database: update 2006. Nucleic Acids Research 2006, 34:D590–598.

65. Loyfer N, Rosenski J, Kaplan T: wgbstools: A computational suite for DNA methylation sequencing data representation, visualization, and analysis. 2024.

66. Shafin K, Pesout T, Chang P-C, Nattestad M, Kolesnikov A, Goel S, Baid G, Kolmogorov M, Eizenga JM, Miga KH, et al: Haplotype-aware variant calling with PEPPER-Margin-DeepVariant enables high accuracy in nanopore long-reads. Nature Methods 2021, 18:1322–1332.

67. Cingolani P, Platts A, Wang LL, Coon M, Nguyen T, Wang L, Land SJ, Lu X, Ruden DM: A program for annotating and predicting the effects of single nucleotide polymorphisms, SnpEff. Fly 2012, 6:80–92.

68. Phan L, Zhang H, Wang Q, Villamarin R, Hefferon T, Ramanathan A, Kattman B: The evolution of dbSNP: 25 years of impact in genomic research. Nucleic Acids Research 2025, 53:D925–D931.

69. Li H: Minimap2: pairwise alignment for nucleotide sequences. Bioinformatics 2018, 34:3094–3100.

70. RepeatMasker [http://www.repeatmasker.org]

71. Tanigawa Y, Dyer ES, Bejerano G: WhichTF is functionally important in your open chromatin data? PLoS computational biology 2022, 18:e1010378.

72. O’Leary NA, Wright MW, Brister JR, Ciufo S, Haddad D, McVeigh R, Rajput B, Robbertse B, Smith-White B, Ako-Adjei D, et al: Reference sequence (RefSeq) database at NCBI: current status, taxonomic expansion, and functional annotation. Nucleic Acids Research 2016, 44:D733–745.

73. Subramanian A, Tamayo P, Mootha VK, Mukherjee S, Ebert BL, Gillette MA, Paulovich A, Pomeroy SL, Golub TR, Lander ES, Mesirov JP: Gene set enrichment analysis: A knowledge-based approach for interpreting genome-wide expression profiles. Proceedings of the National Academy of Sciences 2005, 102:15545–15550.

74. Liberzon A, Subramanian A, Pinchback R, Thorvaldsdóttir H, Tamayo P, Mesirov JP: Molecular signatures database (MSigDB) 3.0. Bioinformatics 2011, 27:1739–1740.

75. Chen S, Zhou Y, Chen Y, Gu J: fastp: an ultra-fast all-in-one FASTQ preprocessor. Bioinformatics 2018, 34:i884–i890.

76. Dobin A, Davis CA, Schlesinger F, Drenkow J, Zaleski C, Jha S, Batut P, Chaisson M, Gingeras TR: STAR: ultrafast universal RNA-seq aligner. Bioinformatics 2013, 29:15–21.

77. Liao Y, Smyth GK, Shi W: featureCounts: an efficient general purpose program for assigning sequence reads to genomic features. *Bioinformatics (Oxford*, England) 2014, 30:923–930.

78. Bendall ML, Mulder Md, Iñiguez LP, Lecanda-Sánchez A, Pérez-Losada M, Ostrowski MA, Jones RB, Mulder LCF, Reyes-Terán G, Crandall KA, et al: Telescope: Characterization of the retrotranscriptome by accurate estimation of transposable element expression. PLOS Computational Biology 2019, 15:e1006453.

79. Love MI, Huber W, Anders S: Moderated estimation of fold change and dispersion for RNA-seq data with DESeq2. Genome Biology 2014, 15:550.

80. Kramer NE, Davis ES, Wenger CD, Deoudes EM, Parker SM, Love MI, Phanstiel DH: Plotgardener: cultivating precise multi-panel figures in R. Bioinformatics 2022, 38:2042–2045.

81. Ritchie ME, Phipson B, Wu D, Hu Y, Law CW, Shi W, Smyth GK: limma powers differential expression analyses for RNA-sequencing and microarray studies. Nucleic Acids Research 2015, 43:e47.

